# CircFOXK2 Promotes Tumor Growth and Metastasis of Pancreatic Ductal Adenocarcinoma via Complexing with RNA Binding Proteins and Sponging MiR-942

**DOI:** 10.1101/792101

**Authors:** Chi Hin Wong, Ut Kei Lou, Youjia Li, Stephen Lam Chan, Joanna Hung-Man Tong, Ka-Fai To, Yangchao Chen

**Affiliations:** School of Biomedical Sciences, Faculty of Medicine, The Chinese University of Hong Kong, Shatin, NT, Hong Kong; Department of Clinical Oncology, Prince of Wales Hospital, The Chinese University of Hong Kong, Shatin, Hong Kong; Department of Anatomical and Cellular Pathology, Prince of Wales Hospital, The Chinese University of Hong Kong, Shatin, Hong Kong; Shenzhen Research Institute, The Chinese University of Hong Kong, Shenzhen 518087, China

**Author notes:** Contact Information Yangchao Chen, PhD, School of Biomedical Sciences, Faculty of Medicine, Chinese University of Hong Kong, Shatin, Hong Kong.

**Keywords:** pancreatic cancer, circular RNA, RNA binding protein, microRNA sponge

## Abstract

**Objective:** Circular RNA (circRNA) is a novel class of non-coding RNAs that regulate gene expression. However, the role of circRNAs in pancreatic ductal adenocarcinoma (PDAC) is largely unknown.

**Design:** We performed circRNA sequencing of non-tumor HPDE and PDAC cells. We investigated the functions of circFOXK2 in PDAC by gain-of-function and loss-of-function assays. Bioinformatics analysis, luciferase assay and microRNA pulldown assays were performed to identify circFOXK2 interacting-miRNAs. To further investigate the mechanism, we performed circRNA-pulldown and mass spectrometry to identify circFOXK2-interacting proteins in PDAC.

**Results:** We identified 169 differentially expressed circRNAs in PDAC cells. We validated that one of the circRNAs circFOXK2 was significantly up-regulated in PDAC cells and in 63 % of primary tumor (53 out of 84). Gain-of-function and loss-of-function assays demonstrated that circFOXK2 promoted PDAC cell growth, migration and invasion. CircFOXK2 was also involved in cell cycle progression and apoptosis. circFOXK2 functioned as sponge for miR-942, and in turn promoted the expression of miR-942 targets ANK1, GDNF and PAX6. Furthermore, circFOXK2 interacted with 94 proteins, which were involved in cell adhesion and mRNA splicing. Among these circFOXK2-interacting proteins, YBX1 and hnRNPK were validated by RNA immunoprecipitation. Importantly, circFOKX2 interacted with YBX1 and hnRNPK targets NUF2 and PDXK in PDAC cells. Knockdown of circFOXK2 reduced the binding of YBX1 and hnRNPK to NUF2 and PDXK, and in turn decreased their expressions in PDAC cells.

**Conclusion:** We identified that circFOXK2 promoted PDAC cells growth and metastasis. Also, circFOXK2 complexed with YBX1 and hnRNPK to promote the expressions of oncogenic proteins.

**Significance of this study:** What is already known on this subject?

- Differentially expressed circRNAs are involved in carcinogenesis of many cancers.
- CircRNAs function as microRNA sponges to regulate gene expression.
- The roles of circRNAs in PDAC progression is largely unknown.

What are the new findings?

- circFOXK2 is upregulated in PDAC primary tumors.
- circFOXK2 promotes PDAC tumor growth and liver metastasis.
- circFOXK2 functions as sponges for miR-942 to promote the expressions of oncogenic ANK1, GDNF and PAX6.
- circFOXK2 complexes with YBX1 and hnRNPK to promote the expressions of oncogenic proteins in PDAC.

How might it impact on clinical practice in the foreseeable future?

- circFOXK2 upregulation in PDAC may function as a novel biomarker for diagnosis.
- circFOXK2 may be a novel therapeutic target in treating PDAC.

## Introduction

Pancreatic ductal adenocarcinoma (PDAC) is the fourth leading cause of cancer death in the USA and Europe^1,2^. Surgery, chemotherapy and radiotherapy are commonly used to treat PDAC. Unfortunately, they can only extend survival for several months^3,4^. Chemotherapy drug gemcitabine is usually used together with surgery or targeted anti-cancer drug erlotinib. In addition, combining gemcitabine with other chemotherapy drugs such as nab-paclitaxel or human monoclonal antibody such as AGS-1C4D4 can only provide a limited extension of survival in exchange of developing adverse effects including abdominal pain, peripheral neuropathy, and myelosuppression^5,6^. Therefore, novel therapeutic targets are urgently needed for treating PDAC.

Circular RNA (circRNA), which is first discovered in 1976, is a type of non-coding RNA with unknown function previously. circRNA is formed by the linking of 3’ end of exon back to its 5’ end, forming a circular structure. The first functional role of circRNAs was identified in 2013. ciRS-7 (circular RNA sponge for miR-7) interacted and inhibited miR-7 activity in brain^7,8^. Since then, the expression patterns and functional roles of circRNAs have been studied in many biological processes including cancer progression. F-circSR1 and F-circSR2 promoted cell migration in lung cancer^9^. CircIRAK3 facilitated breast cancer metastasis through sponging miR-3607^10^. circβ-catenin promoted cell proliferation by activating Wnt pathway in liver cancer^11^. However, the importance of circRNAs in PDAC development is not fully understood.

In this study, we identified 169 differentially expressed circRNAs in PDAC cells by circRNA sequencing. We further validated that one of the circRNAs circFOXK2 was significantly up-regulated in both PDAC cells and tumors. circFOXK2 promoted cell growth, clonogenic ability, migration, invasion and liver metastasis in PDAC. Also, circFOXK2 functioned as microRNA (miRNA) sponge for miR-942, and in turn promoted that expression of Ankyrin 1 (ANK1), Glial cell-derived neurotrophic factor (GDNF), and Pyridoxal Kinase (PAX6). Importantly, circRNA-pull down and mass spectrometry identified 94 circFOXK2-interacting proteins, which were related to many biological processes including cell adhesion, mRNA splicing and cytoskeleton. We demonstrated that circFOXK2 complexed with Y-Box Binding Protein 1 (YBX1) and Heterogeneous Nuclear Ribonucleoprotein K (hnRNPK) to promote the expression of oncogenic proteins NUF2 and PDXK in PDAC.

## Materials and Methods

### Mammalian cell lines and clinical samples

PDAC Cell lines PANC-1, SW1990, CAPAN-2, CFPAC-1, and BxPC-3, and HEK293 cells were obtained from American Type Culture Collection (ATCC). Human pancreatic ductal epithelial (HPDE) cell line was a gifted from Dr Ming-Sound Tsao (University Health Network, Ontario Cancer Institute and Princess Margaret Hospital Site, Toronto)^12^. All cell lines were cultured under the condition as previously described^13^. 84 pairs of PDAC tumor and adjacent non-tumor tissues were obtained from patients who underwent pancreatic resection at the Prince of Wales Hospital, Hong Kong. The study was carried out with the approval of the Joint CUHK-NTEC Clinical Research Ethics Committee. All specimens were fixed and embedded into paraffin

### circRNA sequencing and identification

circRNA sequencing was performed to analyze the expression pattern of circRNAs in HPDE, PANC-1 and SW1990 cells. The total RNA extracted from cells were depleted from ribosomal RNA to linear RNA by RNase R. Then RNA was fragmented and was reverse transcripted. After linking with the sequencing adaptor and PCR amplification, a library was constructed for the circRNA sequencing. The raw sequencing reads from each sample were first mapped to the reference human genome using TopHat2^14^. The unmapped reads were extracted and mapped to reference human genome using TopHat-Fusion^15^. Reads were processed into two anchors from both end of the reads. Anchors which were aligned into the same chromosome but in reversed orientation were considered as the potential back-spliced junction reads. Since the common back-spliced junction were GT/AG, GC/AG and AT/AC, back-spliced junction reads with these junctions were extracted. Mapped reads (from TopHat2) and back-spliced reads (from TopHat-Fusion) were used to quantify the abundance of each circRNA candidate, denoted in RPM (Reads Per Million mapped reads). DEGseq was used to compare the expression level of each circRNA between samples.

### *In vivo* subcutaneous injection

Male BALB/c nude mice aged 4 to 6 weeks were acquired from Laboratory Animal Services Centre of the Chinese University of Hong Kong. Animal handling and experimental procedures were approved by the Animal Experimental Ethics Committee of the institute. For tumor growth assay, 6 × 10^5^ cells were resuspended in 1× PBS with 20% matrigel (Corning) and were injected subcutaneously into the right flank of the nude mice (seven mice per group). After tumor formation, tumor growth was monitored every 3-4 days, and the tumor volume was measured by a caliper and calculated by the equation: volume = (Length × width^2^) / 2.

### *In vivo* orthotopic injection

Male BALB/c nude mice aged 4 to 6 weeks were acquired from Laboratory Animal Services Centre of the Chinese University of Hong Kong. Animal handling and experimental procedures were approved by the Animal Experimental Ethics Committee of the institute. For tumor metastasis assay, 5 × 10^5^ cells were resuspended in 1× PBS with 20 % matrigel and were injected orthotopically to the head of the pancreas. Tumor and organs were collected and examined for metastasis.

### miRNA Pull down

miRNA pull down assay was performed by transfecting PANC-1 cells with 100 nM 3’-end biotinylated miRNA mimics. After 24 h transfection, the cells were washed twice with iced PBS, followed by cell lysis using miRNA Pull down lysis buffer. In the miRNA pull down assay, 25 μL Dynabeads MyOne Streptavidin C1 (Thermofisher) was washed three times with miRNA Pull down lysis buffer. Then the beads were blocked with 1 mg/mL yeast tRNA (Thermofisher) and 1 mg/mL BSA at 4°C for 2h with rotation. After that, the beads were washed twice with miRNA Pull down lysis buffer. Biotin-labelled miRNAs were isolated by incubating the beads with 100 μL cell lysate and 100 μL miRNA Pull down lysis buffer at 4°C for 4 h with rotation. The beads were washed twice with miRNA Pull down lysis buffer. Biotin-labelled miRNAs and their interacting RNAs were isolated by TRIZOL Reagent. Detection of miRNA interacting RNAs was performed by RT-qPCR.

### *In vitro* transcription

circRNA overexpression plasmid was digested with XhoI (NEB) to form a linearized template DNA. Then *in vitro* transcription of circRNAs and their parental mRNAs was performed by MEGA script T7 Transcription Kit (ThermoFisher Scientific)^16^. 1 μg template DNA was mixed with 1 μL of ATP solution, 1 μL of CTP solution, 1 μL of GTP solution, 0.9 μL of UTP solution, 0.15 μL of biotin-UTP solution (Epicentre), 1 μL of enzyme mixture and RNase-free water to 10 μL. After *in vitro* transcription at 37°C overnight, template DNA was digested with DNase for 15 mins at 37°C. For circRNA transcription, parental mRNA was digested by 10U RNase R (Epicentre) for 5 h at 37°C. Digestion was terminated by ammonium acetate stop solution and *in vitro* transcripted RNAs were purified by phenol:cholorform:isoamylalcohol solution according to the manufacturer’s protocol.

### RNA Pull down

RNA Pull down assay for investigating RNA-protein interaction was performed by Pierce Magnetic RNA-Protein Pull-Down Kit (ThermoFisher Scientific). 5 μg of the *in vitro* transcripted RNA was heated at 90°C for 2 mins in 1X RNA Capture Buffer, followed by incubation on ice for 2 mins and at room temperature for 30 mins. Then the biotin-labelled RNA was mixed with 75 μL streptavidin magnetic beads, which were pre-washed twice with Tris-Buffer. After incubation at room temperature for 30 mins, the RNA-labelled beads were washed twice with Tris-buffer. 200 μg PANC-1 protein was added to the RNA-labelled beads in 1X Protein-RNA Binding Buffer. The mixture was incubated at 4°C with rotation overnight. Then the RNA-labelled beads with proteins were washed twice with Wash Buffer. RNA-interacting proteins were eluted with 50 μL Elution Buffer by incubation at 37°C for 30 mins.

### Mass spectrometry

Proteins eluted from the RNA pull down assay was resolved by 12% SDS-PAGE gel. When protein bands entered separating gel, electrophoresis was stopped so the bands were “stacked”. The protein bands were stained by staining solution (0.1% Coomassie Blue R250 in 40% ethanol, 10% acetic acid) and were destained by destaining solution. Then the gel was washed with washing solution (50% methanol and 10% acetic acid) to remove potential contaminant. The gel with bands excised and was completely destained (JTBaker). In-gel digestion was performed by reduction and alkylation by 8mM DTT (USB Chemicals) and 40mM iodoacetamide (GE Healthcare), respectively. Protein digestion was performed by overnight trypsin (1ng/µl; Promega) incubation at 37°C. Subsequent tryptic peptides were extracted from the gel with 100 % ACN/3%TFA (JTBaker) and 40% ACN/3%TFA (JTBaker). The peptide extracts were pooled together and SpeedVac dried. Peptides derived from each condition were desalted using C18 StageTips (3M Corp) for LC-MS/MS analysis. Peptides were analyzed Dionex Ultimate3000 nanoRSLC system coupled to Thermo Fisher Orbitrap Fusion Tribid Lumos. Raw mass spectrometry data were processed using Proteome discoverer 2.1 version against Human UniProt FASTA database (July 2017).

### RNA immunoprecipitation

RNA immunoprecipitation was performed by Magna RIP™ RNA-Binding Protein Immunoprecipitation Kit (Millipore) according to the manufacturer’s protocol. Briefly, cells were washed twice with ice-cold PBS, followed by cell lysis using equal volume of RIP lysis buffer. Magnetic beads were washed twice with RIP wash buffer, followed by incubation with 2 μg antibody against hnRNPK (rabbit; proteintech 11426-1-AP); YBX1 (rabbit; proteintech 20339-1-AP); SEPT11 (rabbit; proteintech 14672-1-AP); ILF3 (rabbit; proteintech 19887-1-AP); ASF (rabbit; proteintech 12929-2-AP) and RAB11FIP1 (rabbit; proteintech 16778-1-AP) for 30 mins at room temperature. Immunoprecipitation was performed by incubating cell lysate with magnetic bead-antibody complex overnight at 4°C. Then the beads were washed six times with RIP wash buffer, followed by proteinase K digestion at 37°C for 30 mins. RNA was purified by phenol:chloroform:isoamylalcohol (15593-031, Invitrogen). qRT-PCR was used to analyze the enrichment of RNAs with target proteins.

### Statistical analysis

Statistical analysis was performed by GraphPad Prism 7. Student’s t-test, chi square t-test and Pearson’s correlation were used as appropriate. Data were shown in mean ± SD. Statistically significant was considered when p-value (two-sided) was less than 0.05.

## Supplemental methods are described in Supplemental information

### Results

#### Identification of circRNAs in PDAC cells

To identify the differentially expressed circRNAs in PDAC, we performed circRNA sequencing of non-tumor human pancreatic epithelial cells (HPDE) and PDAC cells PANC-1 and SW1990. In total, 17,158 circRNAs were identified, in which 84% was exonic (supplemental figure 1A). Comparing with non-tumor HPDE cells, we identified 83 up-regulated and 86 down-regulated circRNAs in PANC-1 and SW1990 cells (supplemental figure 1B). To validate the results from circRNA sequencing and the clinical significances of the up-regulated circRNAs, we measured the expressions of selected circRNAs in a panel of PDAC cells and primary tumors (supplemental figure 1C and D). circFOXK2 was significantly up-regulated in PDAC cells (figure 1A). Also, circFOXK2 was dramatically up-regulated in 63 % of primary tumor (53 out of 84) (figure 1B). Collectively, circFOXK2 was significantly up-regulated in PDAC cells and primary tumors.

**Figure 1.**
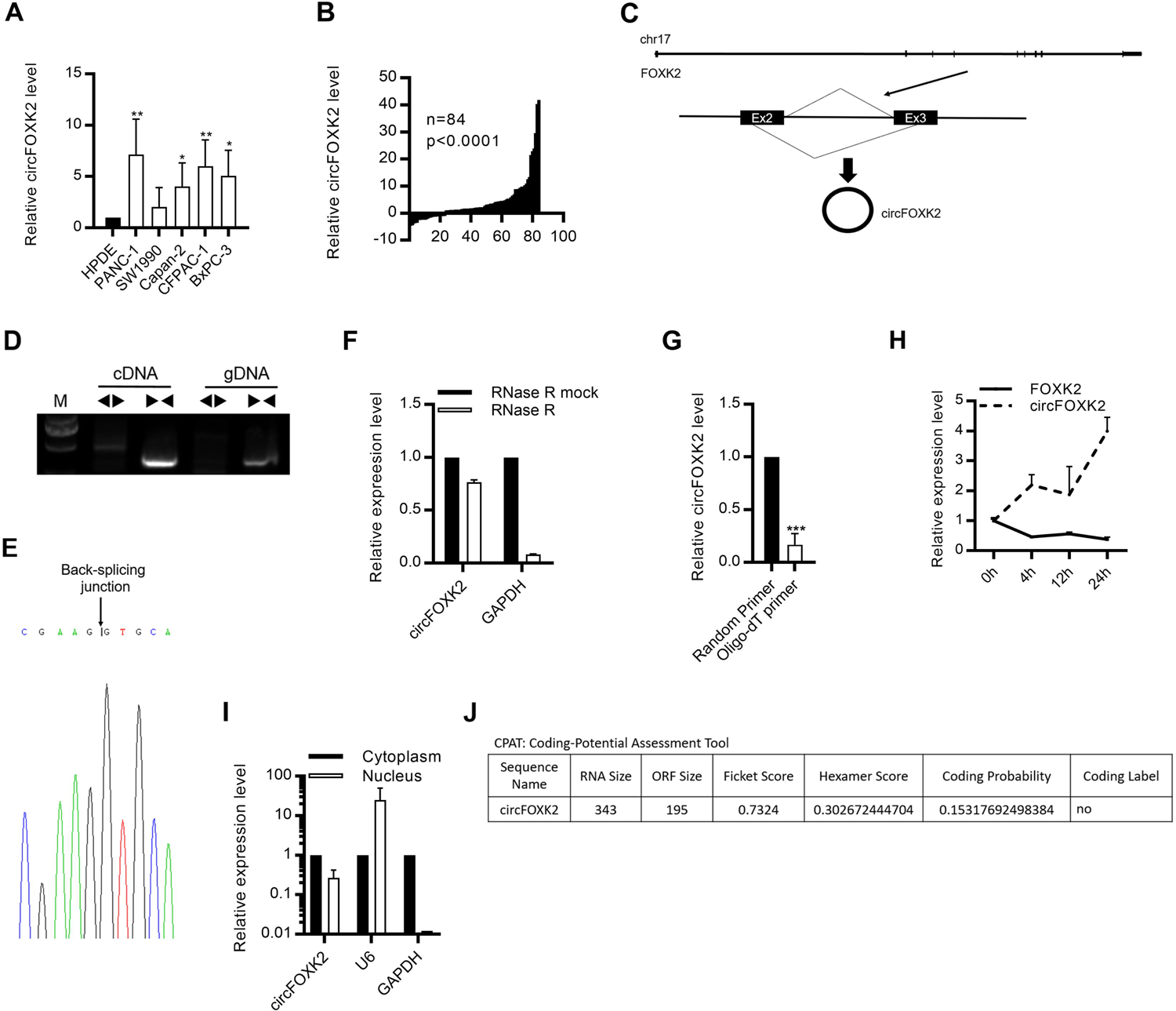
Characterization of circFOXK2 in PDAC. (A) The expression level of circFOXK2 in PDAC cells. The expression level was compared to non-tumor HPDE cells. (B) The expression level of circFOXK2 in PDAC primary tumor samples. The expression level of circFOXK2 in tumor sample was compared to respective adjacent non-tumor tissue. (C) Genomic location of circFOXK2. circFOXK2 was formed by the back-splicing of exon 2 and 3 of FOXK2. (D) PCR of gDNA and cDNA using divergent and convergent primers of circFOXK2. (E) circFOXK2 was amplified using divergent primer followed by sanger sequencing of back-splicing junction. (F) Expression level of circFOXK2 and GAPDH after RNase R treatment of total RNAs from PANC-1 cells. (G) circFOXK2 expression level after reverse transcription with random or oligo-dT primers using total RNAs from PANC-1 cells. (H) Expression level of circFOXK2 and FOXK2 after actinomycin D treatment for 0h, 4h, 12h and 24h in PANC-1 cells. (I) Cellular localization of circFOXK2 in PANC-1 cells. nuclear and cytoplasmic fraction was separated followed by RNA extraction. circFOXK2, U6 and GAPDH levels were analyzed by qRT-PCR. (J) Analysis of coding potential of circFOXK2 by Coding Potential Assessment Tool. Data are from at least three independent experiments Mean ± SD (***p<0.001)

#### Characterization of circFOXK2

We next validated circFOXK2 through examining its physical circular structure. circFOXK2 is formed by the back-splicing of exon 2 and 3 of Forkhead box protein K2 (FOXK2) (figure 1C). Outward-facing divergent primer and inward-facing convergent primer were designed to validate the formation of circFOXK2. Both divergent and convergent primer amplified a product of expected size from complementary DNA (cDNA), while only convergent amplified a product from genomic DNA (gDNA) (figure 1D). The presence of back-splicing junction was validated by Sanger sequencing (figure 1E). Also, we found that circFOXK2 was resistant to the digestion by RNase R which specifically degraded linear RNAs but not the circRNAs (figure 1F). The reduced efficiency of reverse-transcription by oligo-dT primers due to the lack of polyA tail also demonstrated the circularity of circFOXK2 (figure 1G). These results suggested the formation of circFOXK2 was not due to genomic rearrangement. Because of the circular structure, we found that circFOXK2 was more stable than FOXK2 (figure 1H). Localization of circFOXK2 was examined by measuring its level in cytoplasm and nucleus. circFOXK2 was enriched in the cytoplasm in PDAC cells (figure 1I). Coding potential analysis suggested that were lack of protein coding ability (figure 1J). Our results validated the circularity of circFOXK2.

#### circFOXK2 promotes cell growth and invasion *in vitro*

To elucidate the functions of circFOXK2 in PDAC, we employed small interfering RNA (siRNA) which specifically targeted the back-splicing junction of circFOXK2; without altering the expression of its parental gene (supplemental figure 2A). Knockdown of circFOXK2 inhibited cell growth and clonogenic ability of PDAC cells (figure 2A-C). The reduced cell growth by knockdown of circFOXK2 was due to cell cycle arrest in G0/G1 phase and induction of apoptosis (figure 2D). Annexin V staining, cleavage of PARP and caspase 3 and decrease in Bcl-2 level also demonstrated the induction of apoptosis after knockdown of circFOXK2 in PDAC cells (figure 2E and F). Also, knockdown of circFOXK2 inhibited cell migration and invasion (figure 2G and H). Furthermore, we constructed stable knockdown CFPAC-1 cells using sh-circFOXK2 lentiviral system. Stable knockdown of circFOXK2 inhibited cell growth, clonogenic ability and invasiveness (supplemental figure 2B-E). On the other hand, overexpression of circFOXK2 in HPDE cells significantly promoted cell growth, clonogenic ability, migration and invasion. (supplemental figure 3). Collectively, our results demonstrated the importance of circFOXK2 in promoting PDAC cell growth and invasion.

**Figure 2.**
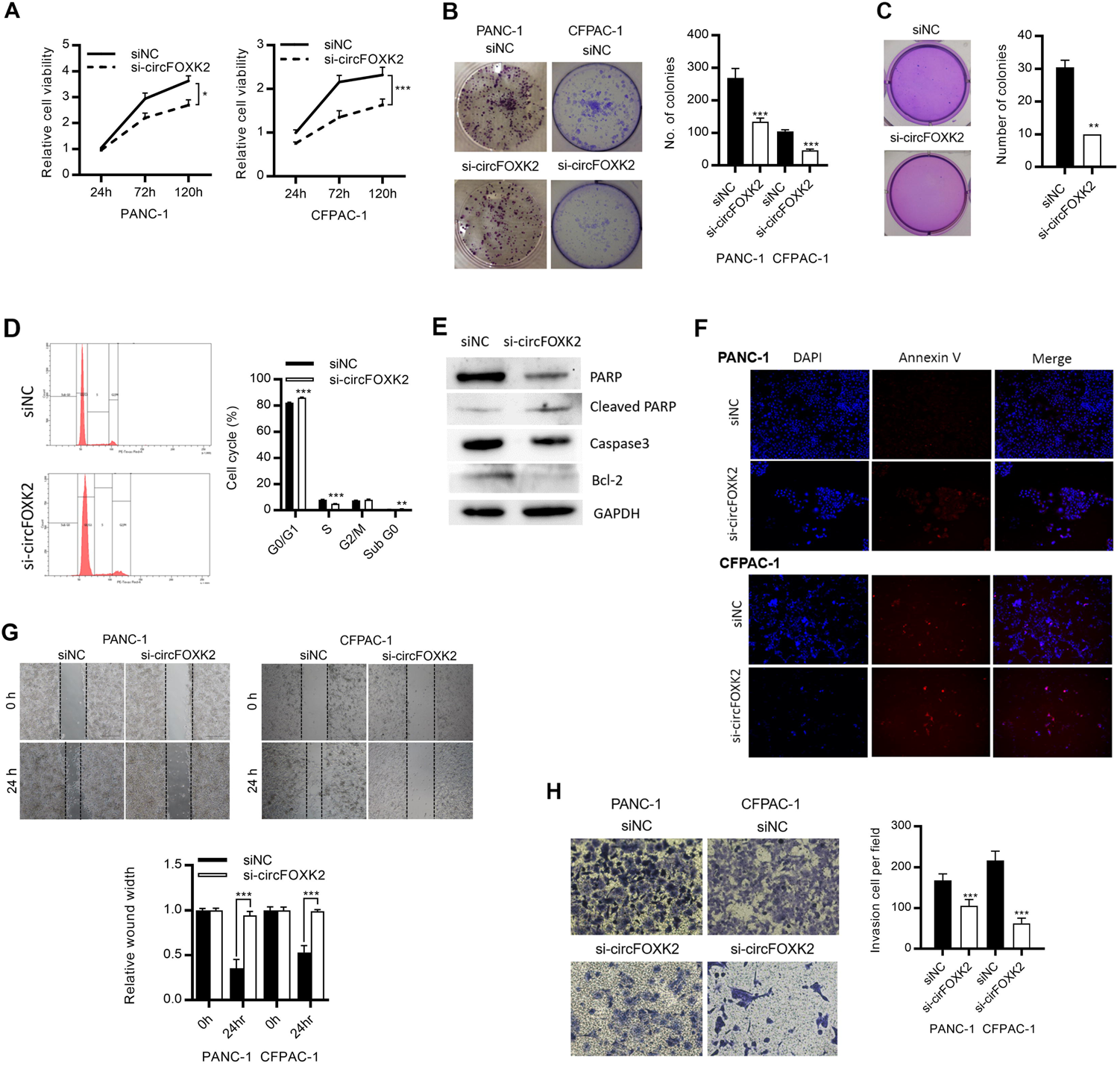
circFOXK2 promotes cell growth, migration and invasion in PDAC cells. (A) Cell growth analyzed by MTT assay after knockdown of circFOXK2 by siRNA for 24, 72 and 120 h in PDAC cells. (B) Anchorage-dependent colony formation after knockdown of circFOXK2 in PDAC cells. Cells were stained with crystal violet. (C) Anchorage-independent colony formation after knockdown of circFOXK2 in PANC-1 cells. Cells were stained with crystal violet. (D) Analysis of cell cycle distribution by flow cytometry after knockdown of circFOXK2 in CFPAC-1 cells for 72 h. (E) Western blot of apoptosis marker PARP, Bcl-2, Caspase 3 after knockdown of circFOXK2 in CFPAC-1 cells. (F) Cell apoptosis assessed by annexin V staining (red) after knockdown of circFOXK2in PADC cells for 72 h.(G-H) (G) Wound healing cell migration assay and (H) Trans-well cell invasion assay after knockdown of circFOXK2 in PDAC cells. Cells in invasion assay were stained by crystal violet. Data are from at least three independent experiments Mean ± SD (*p<0.05; **p<0.01; ***p<0.001)

#### circFOXK2 promotes tumor growth and metastasis *in vivo*

Knockdown of circFOXK2 was also employed for *in vivo* study. Xenograft mice model was generated by subcutaneous injection of sh-circFOXK2 CFPAC-1 cells. Knockdown of circFOXK2 by shRNA significantly inhibited tumor growth (figure 3A-D). Since *in vitro* data revealed that circFOXK2 promoted PDAC cell migration and invasion, we constructed PDAC metastatic mice model through orthotopic injection of sh-circFOXK2 CFPAC-1 cells to the pancreas. Knockdown of circFOXK2 inhibited the PDAC metastasis to liver (figure 3E). Our results found that that circFOXK2 played critical roles in promoting PDAC growth and liver metastasis.

**Figure 3.**
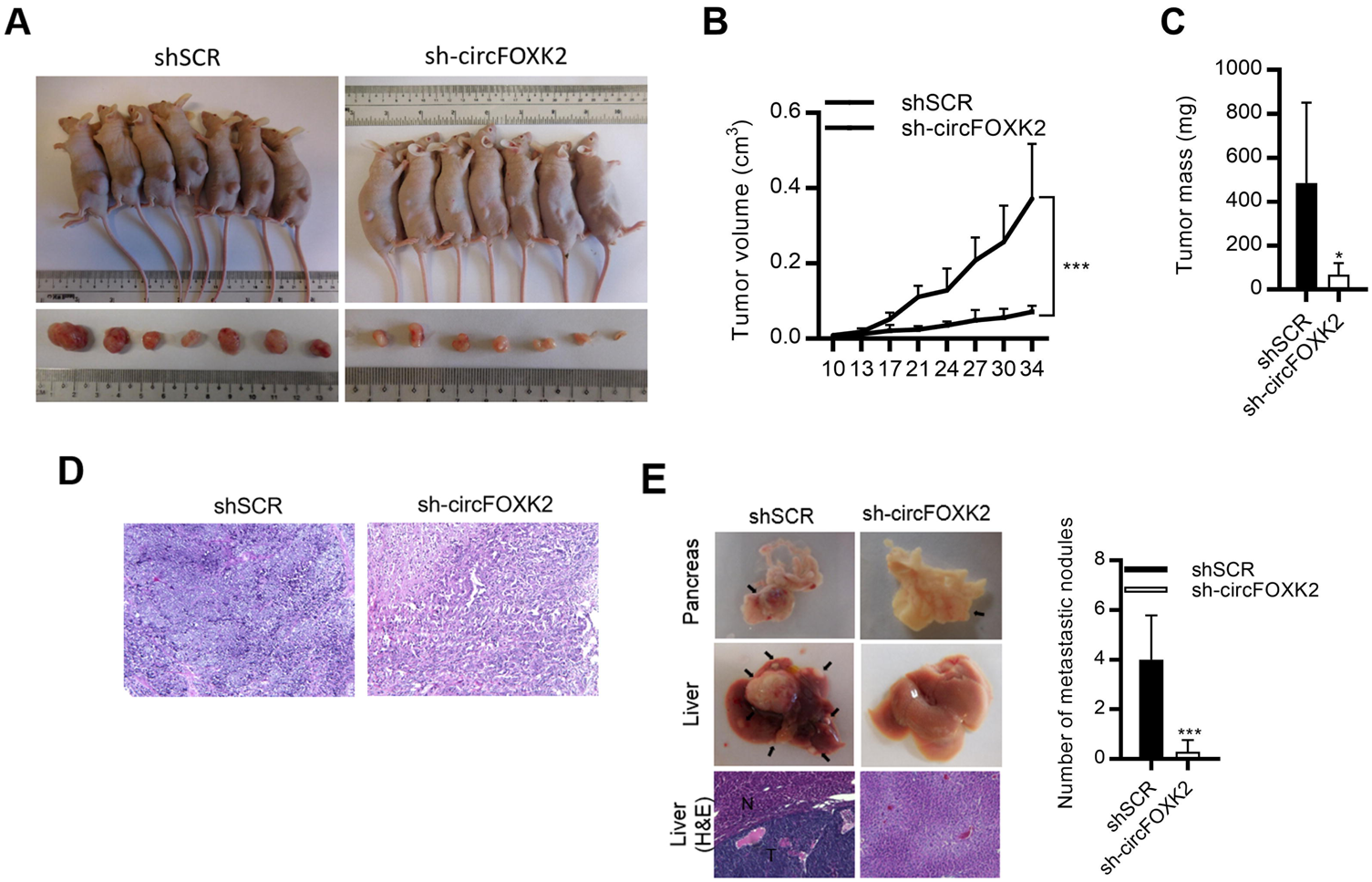
circFOXK2 promotes tumor growth and liver metastasis in PDAC cells. (A) Photographs of mice xenograft at day 34 after knockdown of circFOXK2 in CFPAC-1 cells. (B) Tumor volume of xenograft after knockdown of circFOXK2. (C) Tumor volume of xenograft after knockdown of circFOXK2. (D) H&E staining of mice xenograft after knockdown of circFOXK2. (E) Analysis of tumor metastasis to liver after knockdown of circFOXK2 in CFPAC-1 cells. Data are from at least three independent experiments Mean ± SD (*p<0.05; ***p<0.001)

#### Identification of circFOXK2 interacting miRNAs

We next investigated the detailed mechanism of circFOXK2 in PDAC. Since many cytoplasmic circRNAs serves as “microRNA sponge” in regulating cancer progression, including liver, lung and breast cancer^9-11^, we investigated whether circFOXK2 could function as an microRNA sponge. First, we performed bioinformatics analysis by TargetScan^18^ to reveal potential microRNA binding sites on circFOXK2. To validate the circFOXK2-microRNA interaction, luciferase reporter assay using circFOXK2-luciferase reporter and miRNA mimics was performed (figure 4A). We found that there was a reduction in luciferase activity for miR-942 mimics (figure 4B). Mutating the miR-942 binding sites restored the luciferase activity (figure 4C). circFOXK2-microRNA interaction was further validated by microRNA pull down. Biotin-labelled miR-942 mimics could significantly enriched circFOXK2 in PANC-1 cells (figure 4D). These results suggested the interaction between circFOXK2 and miR-942 in PDAC cells.

**Figure 4.**
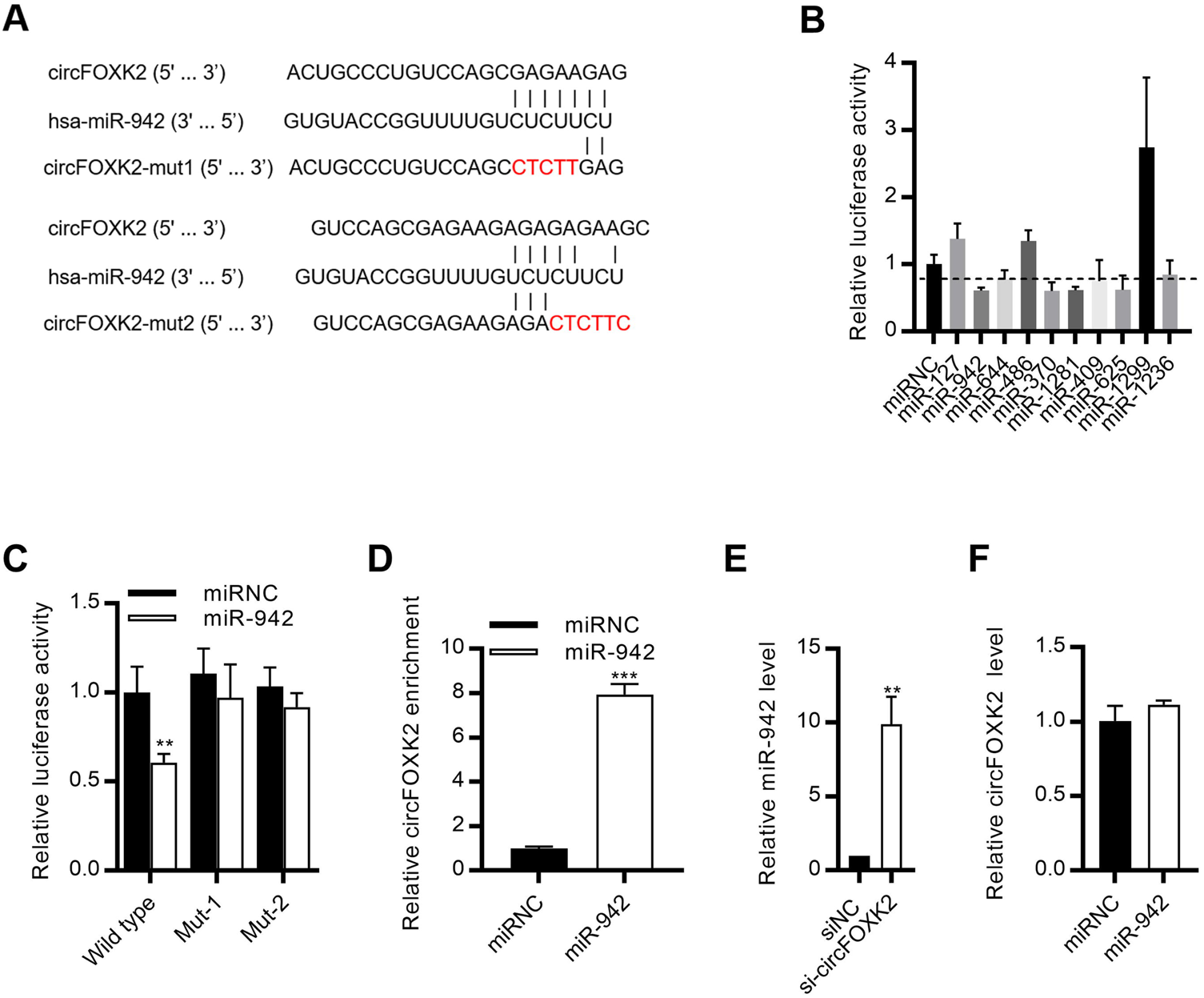
circFOXK2 functions as miR-942 sponge. (A) Bioinformatic analysis of potential miRNA binding sites on circFOXK2. Potential binding sites were mutated for luciferase assay. (B) Luciferase reporter assay for luciferase activity of circFOXK2-reporter in HEK293 cells co-transfected with miRNA mimics. (C) Luciferase reporter assay for luciferase activity of circFOXK2-reporter with mutated miR-942 binding sites in HEK293 cells co-transfected with miR-942 mimics. (D) miRNAs pull down assay in PANC-1 cells co-transfected with biotin-labelled miR-942 mimics. The interaction between circFOXK2 and miR-942 was analyzed by qRT-PCR. (E) miR-942 expression level after knockdown of circFOXK2 in PANC-1 cells. (F) circFOXK2 expression level after transfecting miR-942 mimics in PANC-1 cells. Data are from at least three independent experiments Mean ± SD (**p<0.01; ***p<0.001)

In order to study the functions of circFOXK2-miR-942 interaction in PDAC, we first knocked down circFOXK2. Knockdown of circFOXK2 increased miR-942 expression level (figure 4E). However, transfecting miR-942 mimics did not alter the expression of circFOXK2, but down-regulated the targets of miR-942: ANK1, GDNF and PAX6 (figure 4F and supplemental figure 4). These results suggested that circFOXK2 functioned as sponge for miR-942 to inhibit the activity of miR-942 in PDAC. As the role of miR-942 in PDAC is unclear, we attempted to study the importance of miR-942 in PDAC. We found that miR-942 was frequently down-regulated in PDAC cells and tumors (supplemental figure 5A-C). In addition, we found that transfecting miR-942 mimics inhibited cell growth and clonogenic ability in PDAC cells (supplemental figure 5D and E). Our results found that miR-942 promoted PDAC cell growth.

Since circFOXK2 functioned as miRNA sponge for miR-942, the expression level of ANK1, GDNF and PAX6 were measured after knockdown or overexpression of circFOXK2. Knockdown of circFOXK2 decreased the expression levels of ANK1, GDNF and PAX6 in PDAC cells, while overexpression of circFOXK2 promoted the expression of ANK1, GDNF and PAX6 in HPDE cells (figure 5A and B and supplemental figure 6). In addition, in the mice xenograft with circFOXK2 knockdown, ANK1, GDNF and PAX6 were down-regulated (figure 5C). Analysis of PDAC primary tumors also revealed the positive correlation between circFOXK2, ANK1 and PAX6 levels (figure 5D). Collectively, our results suggested that circFOXK2 inhibited miR-942, and in turn promoted the expression of ANK1, GDNF and PAX6 in PDAC.

**Figure 5.**
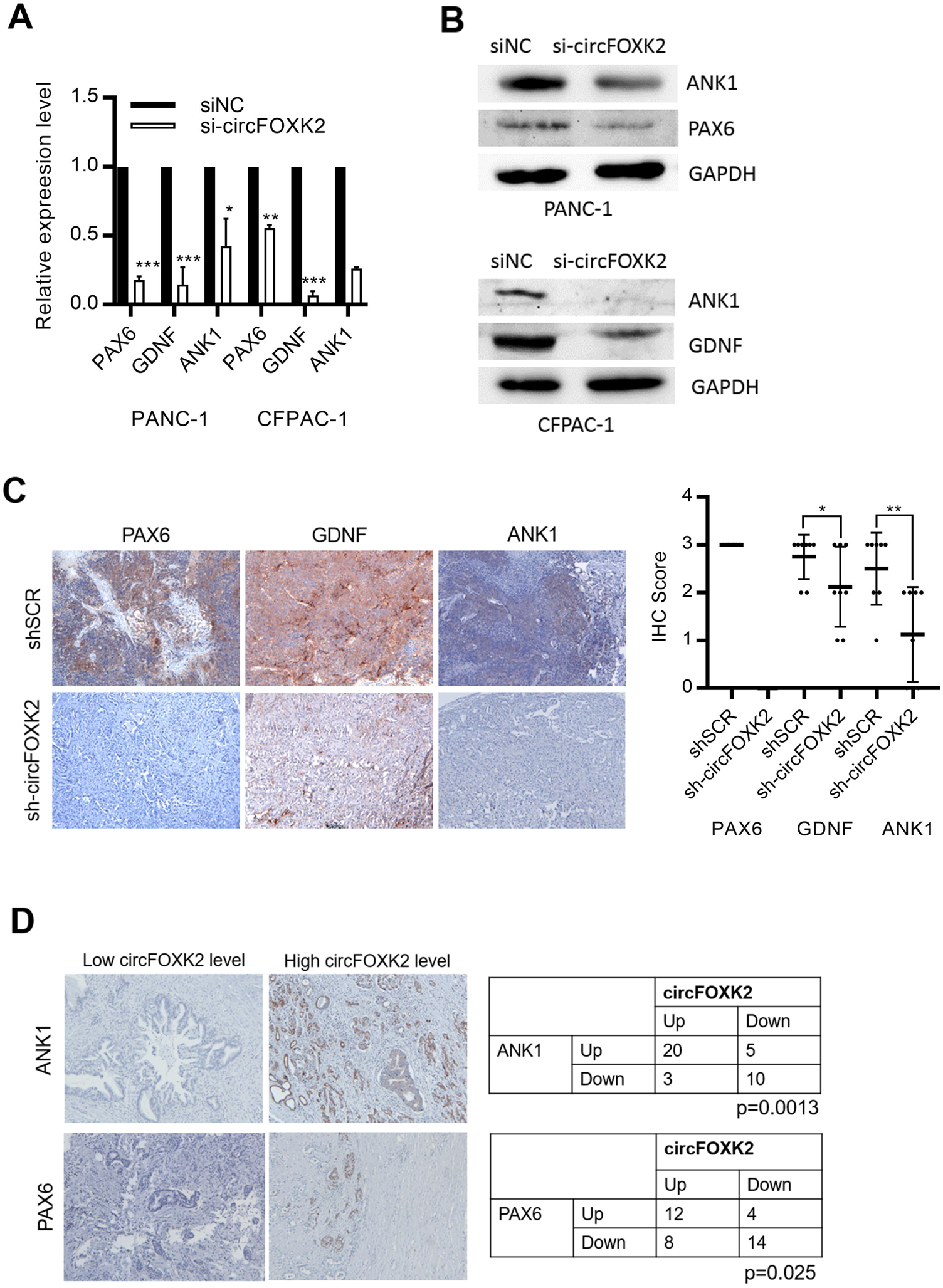
circFOXK2 promotes the expression of miR-942 targets in PDAC. (A-B) qRT-PCR analysis of (A) mRNA and Western blots analysis of (B) protein levels of ANK1, GDNF and PAX6 after knockdown of circFOXK2 in PDAC cells. (C) Immunohistochemistry staining of ANK1, GDNF and PAX6 in mice xenograft after knockdown of circFOXK2. (D) Correlation between circFOXK2 and ANK1 and PAX6 levels in PDAC tumors. Data are from at least three independent experiments Mean ± SD (*p<0.05; **p<0.01; ***p<0.001)

#### Identification of circFOXK2 interacting proteins

Many studies have demonstrated that functioning as microRNA sponges was an important role of circRNAs. However, studies on whether circRNAs could bind to protein is still limited. Importantly, how circRNA-protein interaction contributes to the regulation of gene expression is largely unknown. Therefore, we next studied the importance of circRNA-protein interaction in PDAC. We *in vitro* transcripted circFOXK2, which functioned as a probe to pull-down circFOXK2-interacting proteins. In order to eliminate the “false positive” of the pull-down proteins, we constructed additional two negative controls: 1) RNA probe with identical sequence as circFOXK2, without the formation of circRNAs; 2) RNA probe with similar secondary structure as circRNA, which is circGFP (figure 6A). Therefore, the proteins pull-down by circFOXK2 were due to its unique sequence and secondary structure. Mass-spectrometry analysis identified 94 circFOXK2-interacting proteins in PANC-1 cells (figure 6B). In addition, protein-protein interactions (PPI) of the circRNA-interacting proteins were analyzed in the STRING database^19^. 101 PPI were observed for circFOXK2-interacting proteins (supplemental figure 7A). Gene ontology analysis also revealed that the circFOXK2-interacting proteins play important roles in many biological processes, including cell adhesion, mRNA splicing and cytoskeleton (supplemental figure 7B). Our results suggested that circFOXK2 interacted with multiple proteins with critical roles in PDAC.

### circFOXK2 promotes the expression of oncogenic proteins via complexing YBX1 and hnRNPK

RNA immunoprecipitation assay was performed in PDAC cells to validate the results from mass spectrometry analysis. circFOXK2 was significantly enriched by YBX1 and hnRNPK (figure 6C). Since YBX1 and hnRNPK form a complex in regulating gene expression^19^, we hypothesized that circFOXK2 interacted with YBX1 and hnRNPK complex to promote PDAC development. Publicly available dataset of knockdown of YBX1 was first employed to identify genes that were consistently regulated by YBX1 in multiple cancers (supplemental figure 8A). RNA immunoprecipitation assay demonstrated that Coronin 1C (CORO1C), NUF2 Component Of NDC80 Kinetochore Complex (NUF2), Pyridoxal Kinase (PDXK) and Protein Phosphatase 2 Regulatory Subunit B gamma (PPP2R2C) were enriched by YBX1 and hnRNPK, indicating they are the direct targets of YBX1 and hnRNPK complex (supplemental figure 8B). Consistently, knockdown of YBX1 significantly down-regulated these targets in CFPAC-1 cells (supplemental figure 8C). Analysis of TCGA dataset also revealed the up-regulation of PPP2R2C, PDXK and NUF2, and their positive correlations to YBX1 expression in PDAC (supplemental figure 8D-F). Importantly, NUF2 and PDXK were sensitive to the knockdown of circFOXK2 and were significantly enriched by circFOXK2 pull-down (figure 6D and E). Knockdown of circFOXK2 decreased the interaction of YBX1 and hnRNPK to NUF2 and PDXK (figure 6F and G). Analysis of our PDAC primary tumors revealed the up-regulation of NUF2, and its positive correlation to circFOXK2 expression (figure 7H). Collectively, our results suggested that circFOXK2, YBX1 and hnRNPK complex interacted and promoted the expression of oncogenic proteins NUF2 and PDXK in PDAC.

**Figure 6.**
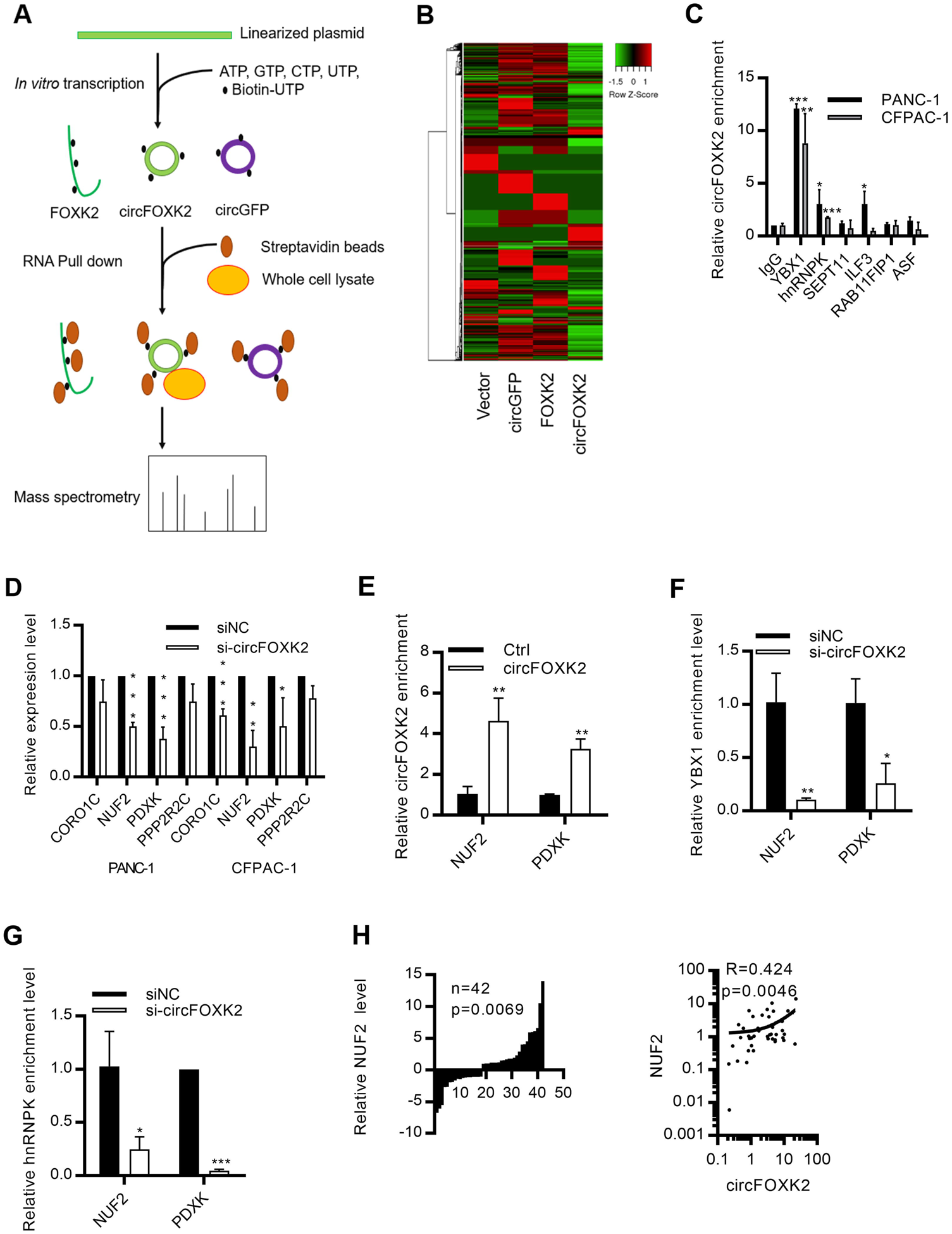
Identification of circFOXK2-interacting proteins in PDAC. (A) Schematic diagram showing the process of circFOXK2-pull down. circFOXK2, FOXK2 and circGFP probes were generated by *in vitro* transcription with biotin-UTP. The biotin-labelled probes were used to pull down interacting proteins in PANC-1 cells. The pull-down proteins were identified by mass spectrometry. (B) Mass spectrometry analysis of circFOXK2-interacting proteins in PANC-1 cells. (C) RNA immunoprecipitation for YBX1, hnRNPK, SEPT11, ILF3, RAB11FIP1 and ASF in PDAC cells. The interactions between circFOXK2 and YBX1, hnRNPK, SEPT11, ILF3, RABIIFIP1 and ASF were analyzed by qRT-PCR. (D) Expression levels of YBX1 and hnRNPK targets CORO1C, NUF2, PDXK and PPP2R2C after knockdown of circFOXK2 in PDAC cells. (E) circFOXK2-pull down assay in PANC-1 cells using biotin-labelled circFOXK2 probe. The interaction between circFOXK2, NUF2 and PDXK was analyzed by qRT-PCR. (F-G) RNA immunoprecipitation for (F) YBX1 and (G) hnRNPK after knockdown of circFOXK2 in PANC-1 cells. The NUF2 and PDXK enrichment were analyzed by qRT-PCR. (H) Expression levels of NUF2 and its correlations to circFOXK2 level in PDAC tumors. Data are from at least three independent experiments Mean ± SD (*p<0.05; **p<0.01; ***p<0.001)

**Figure 7.**
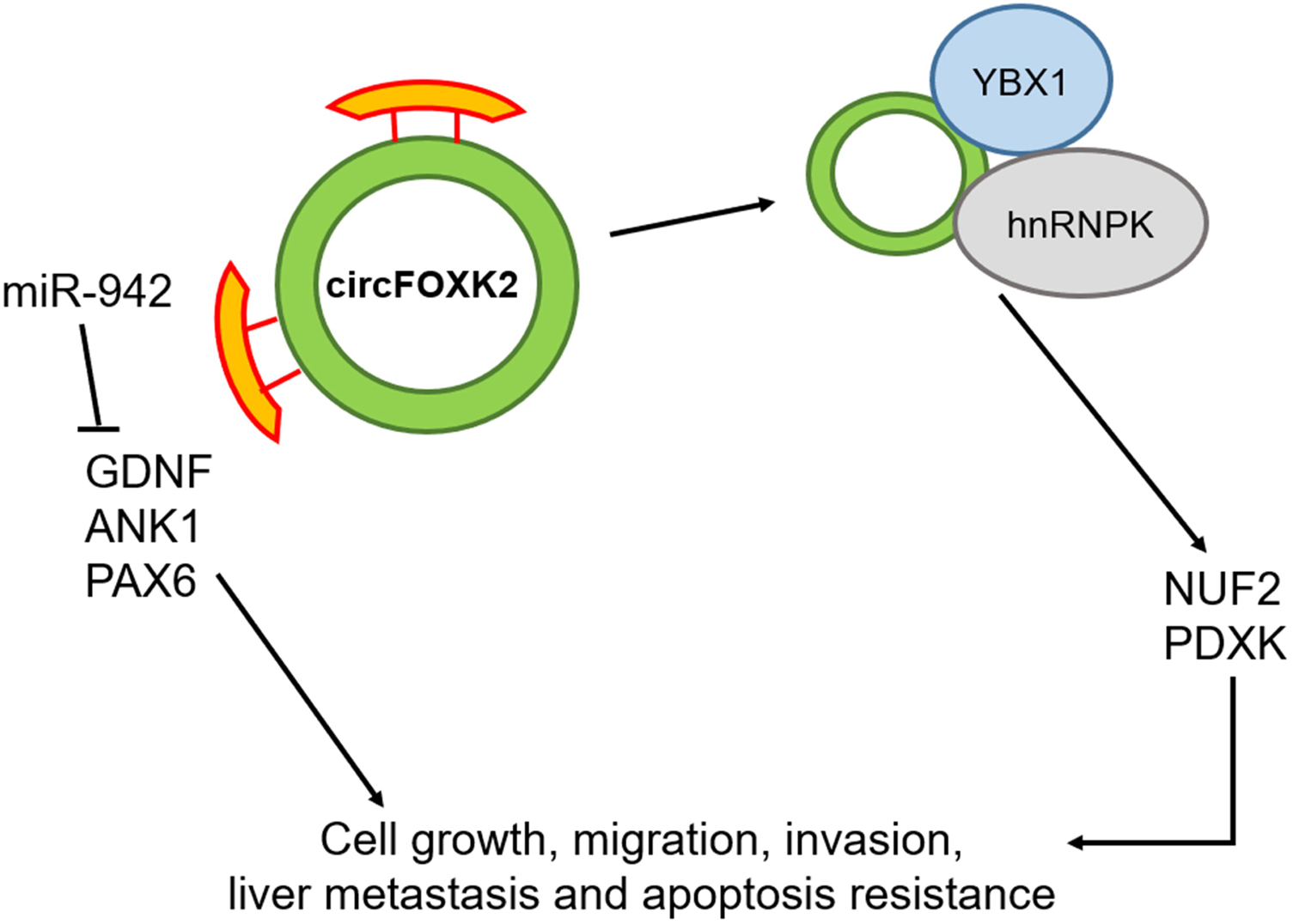
Schematic diagram describing the role of circFOXK2 in the development and progression of PDAC. circFOXK2 functioned as sponge for miR-942, and in turn promoted the expression of ANK1, GDNF and PAX6. In addition, circFOXK2 interacted with YBX1 and hnRNPK to promote the expression of NUF2 and PDXK. These pathways favored cell growth, migration, invasion and liver metastasis in PDAC.

### Discussion

In the current study, we profiled circRNAs expression in non-tumor HPDE and PDAC cells by circRNA sequencing. We identified 169 circRNAs which were differentially expressed in PDAC cells. Further validation experiment demonstrated that circFOXK2 was up-regulated in PDAC cells and tumors. Loss-of-function by both siRNA and shRNA and gain-of-function studies suggested that circFOXK2 played important roles in PDAC by enhancing cell growth, migration, invasion and metastasis *in vitro* and *in vivo*. Also, circFOXK2 functioned as miRNA sponges and interacted with multiple proteins to regulate gene expression.

Many studies have illustrated the importance of circRNAs in cancer progression, including liver, lung and breast cancer^9-11^. Previous circRNA profiling in 20 PDAC tissues identified 289 dysregulated circRNAs^20^. Another study using 6 PDAC tissues discovered 351 differentially expressed circRNAs in PDAC^21^. In general, these differentially expressed circRNAs were involved in protein localization, RNA binding, chromatin modification and protein binding^20^. circRNA_100782 functioned as miR-124 sponge to promote PDAC cell proliferation via IL6-STAT3 pathway^22^. hsa_circ_0006215 promoted migration via inhibiting miRIZI 378aIZI 3p in BxPC-3 cells^23^. circ-PDE8A stimulated MACC1 expression by sponging miR-338 to promote cell invasion^24^. However, the detailed roles of circRNAs in PDAC are still very unclear. Therefore, our study is the first to perform circRNA sequencing to identify differentially expressed circRNAs in PDAC. We identified 83 up-regulated and 86 down-regulated circRNAs in PDAC cells. In addition, we demonstrated that circFOXK2 was up-regulated in PDAC cells and tumors. Importantly, we demonstrated that circFOXK2 played important roles in PDAC cells through promoting cell growth, migration, invasion and liver metastasis.

Functioning as miRNA sponges were the first identified role of circRNA. CDR1as harbored 63 binding sites for miR-7^7^. The interaction between CDR1as and miR-7 blocked the miR-7 activity. Since that, the functions of circRNAs as a “miRNA sponges” were comprehensively studied in many biological processes. In addition, the importance of circRNA-miRNA interactions was also documented in many cancers, including PDAC. circLARP4 inhibited miR-424-5p and regulated LASTS1 expression in gastric cancer^25^. circITCH absorbed miR-17 and miR-224 to promote p21 and PTEN expression in bladder cancer^26^. ciRS-7 targeted miR-7 to promote EGFR/STAT3 signaling, while circZMYM2 sponged miR335-5p and in turn favored the expression of JMJD2C in PDAC^27,28^. Here, we performed bioinformatics analysis, luciferase assay and miRNA pull down assay to identify circFOXK2-interacting miRNAs in PDAC cells. We observed that circFOXK2 absorbed miR-942, and in turn promoted the expression of ANK1, GDNF and PAX6. Studies found that miR-942 may serve as potential diagnostic marker for pancreatic cancer^29^. However, the roles of miR-942 in PDAC is poorly understood. We found that miR-942 inhibited PDAC cell growth and was the negative regulator of ANK1, GDNF and PAX6, which were involved in PDAC cell growth, invasion and metastasis^30-32^.

Apart from functioning as miRNA sponge, limited studies illustrated the importance of circRNA-protein interaction. circ-Amotl1 interacted with c-myc and promoted c-myc translocation to nucleus in breast cancer^33^. Circ-CTNNB1 interacted with DDX3 to transactivate YY1 and YY1 downstream targets in gastric cancer^34^. Importantly, circRNA-protein interaction has not been documented in PDAC. Here, we developed a novel and highly specific circRNA-pull down assay and identified the interacting proteins of circFOXK2 by mass spectrometry. We identified 94 circFOXK2-interacting proteins in PANC-1 cells. Gene ontology analysis indicated these proteins were associated with multiple biological processes, including cell adhesion, cytoskeleton and mRNA splicing.

We then validated the results from mass spectrometry by RNA immunoprecipitation assay. Notably, we found that YBX1 and hnRNPK interacted with circFOXK2 in both PANC-1 and CFPAC-1 cells. YBX1 functions as RNA binding protein that regulates transcription, mRNA processing and translation. YBX1 was frequently up-regulated and involved in cell invasion and drug resistance development in many cancers^35-37^. For pancreatic cancer, studies reported that YBX1 promoted cell invasion^38^. hnRNPK, as a partner of YBX1 in gene regulation, was also aberrantly expressed in various cancers^39^. hnRNPK regulated cell migration via promoting the expression of GSN in lung adenocarcinoma^40^. The up-regulation of hnRNPK also favored cancer metastasis by promoting the expression of MMP-3^41^. In addition, YBX1 and hnRNPK complex was involved in cancer progression by regulating splicing process^42^. In this study, we found that CORO1C, NUF2, PDXK, PPP2R2C were directly regulated by YBX1 and hnRNPK complex in PDAC cells. Reported studies have demonstrated the growth-promoting role of NUF2 in cancer, including PDAC^43^. Importantly, among these YBX1 and hnRNPK targets, NUF2 and PDXK were also regulated by circFOXK2. Notably, knockdown of circFOXK2 reduced the interaction of YBX1 and hnRNPK to NUF2 and PDXK, suggesting circFOXK2 played an important role in YBX1- and hnRNPK-mediated gene regulation. Collectively, circFOXK2 complexed with YBX1 and hnRNPK to promote the expression of oncogenic proteins NUF2 and PDXK.

In conclusion, we profiled the circRNAs expression in HPDE, PANC-1 and SW1990 cells by circRNA sequencing. circFOXK2 was significantly up-regulated in both PDAC cells and primary tumors. Gain-of-function and loss-of-function studies demonstrated that circFOXK2 played important roles in cell growth, migration, invasion and liver metastasis. We found that circFOXK2 functioned as sponge of miR-942, and in turn promoted the expression of ANK1, GDNF and PAX6 (figure 7). More importantly, we found that circFOXK2 interacted with YBX1 and hnRNPK to promote the expression of oncogenic proteins NUF2 and PDXK (figure 7). These revealed a novel mechanism of circRNA in regulating gene expression by interacting with YBX1-hnRNPK complex.

## Acknowledgments

This work was supported by General Research Fund, Research Grants Council of Hong Kong [4171217 and 14120618]; National Natural Science Foundation of China [81672323]; Direct Grant from CUHK to YC.

Mass spectrometry was performed by The Proteomics and Metabolomics Core Facility, Li Ka Shing Faculty of Medicine, The University of Hong Kong.

## Author Contributions

CHW performed most of the experiments and drafted the manuscript. UKL assisted to obtain data. YL assisted to perform the circRNA sequencing. SLC, JHMT and KFT provided some research materials. YC was the PI of the grant, overlook the whole progress and revised the manuscript.

## Data availability

circRNA sequencing data are available in the NCBI Gene Expression Omnibus under accession number GSE135731. The GEO accession token for reviewer was gdenssignnolzyx. Mass spectrometry data of circRNA pull-down are available via ProteomeXchange with identifier PXD015048. The ProteomeXchange reviewer username and password were; 4RHLjCxH.

## Reference

1. Siegel RL, Miller KD, Jemal A. Cancer statistics, 2018. CA Cancer J Clin 2018; 68: 7–30.

2. Malvezzi M, Bertuccio P, Rosso T, et al. European cancer mortality predictions for the year 2015: does lung cancer have the highest death rate in EU women? Annals of Oncology 2015; 26: 779–786.

3. Cress RD, Yin D, Clarke L, et al. Survival among patients with adenocarcinoma of the pancreas: a population-based study (United States). Cancer Causes Control 2006; 17: 403–409.

4. Springfeld C, Jäger D, Büchler MW, et al. Chemotherapy for pancreatic cancer. Presse Med 2019; 48: e159–e174.

5. Von Hoff DD, Ervin T, Arena FP, et al. Increased survival in pancreatic cancer with nab-paclitaxel plus gemcitabine. The New England Journal of Medicine 2013; 369: 1691–1703.

6. Wolpin BM, O’Reilly EM, Ko YJ, et al. Global, multicenter, randomized, phase II trial of gemcitabine and gemcitabine plus AGS-1C4D4 in patients with previously untreated, metastatic pancreatic cancer. Annals of Oncology 2013; 24: 1792–1801.

7. Memczak S, Jens M, Elefsinioti A, et al. Circular RNAs are a large class of animal RNAs with regulatory potency. Nature 2013; 495: 333–8.

8. Hansen TB, Jensen TI, Clausen BH, et al. Natural RNA circles function as efficient microRNA sponges. Nature 2013; 495: 384–8.

9. Wu K, Liao X, Gong Y, et al. Circular RNA F-circSR derived from SLC34A2-ROS1 fusion gene promotes cell migration in non-small cell lung cancer. Mol Cancer 2019; 18: 98.

10. Wu J, Jiang Z, Chen C, et al. CircIRAK3 sponges miR-3607 to facilitate breast cancer metastasis. Cancer Lett 2018; 430: 179–192.

11. Liang WC, Wong CW, Liang PP, et al. Translation of the circular RNA circβ-catenin promotes liver cancer cell growth through activation of the Wnt pathway. Genome Biol 2019; 20: 84.

12. Ouyang H, Mou Lj, Luk C, et al. Immortal human pancreatic duct epithelial cell lines with near normal genotype and phenotype. Am J Pathol 2000; 157: 1623–31.

13. Li CH, Xiao Z, Tong JH, et al. EZH2 coupled with HOTAIR to silence MicroRNA-34a by the induction of heterochromatin formation in human pancreatic ductal adenocarcinoma. Int J Cancer 2017; 140: 120–9.

14. Kim D, Pertea G, Trapnell C, et al. TopHat2: accurate alignment of transcriptomes in the presence of insertions, deletions and gene fusions. Genome Biol 2013; 14: R36.

15. Kim D, Salzberg SL. TopHat-Fusion: an algorithm for discovery of novel fusion transcripts. Genome Biol 2011; 12: R72.

16. Xu F, Li CH, Wong CH, et al. Genome-Wide Screening and Functional Analysis Identifies Tumor Suppressor Long Noncoding RNAs Epigenetically Silenced in Hepatocellular Carcinoma. Cancer Res 2019; 79:1305–1317.

17. Li CH, To KF, Tong JH, et al. Enhancer of zeste homolog 2 silences microRNA-218 in human pancreatic ductal adenocarcinoma cells by inducing formation of heterochromatin. Gastroenterology 2013; 144: 1086–1097.

18. Grimson A, Farh KK, Johnston WK, et al. MicroRNA Targeting Specificity in Mammals: Determinants beyond Seed Pairing. Molecular Cell 2007; 27: 91–105.

19. Shnyreva M, Schullery DS, Suzuki H, et al. Interaction of two multifunctional proteins. Heterogeneous nuclear ribonucleoprotein K and Y-box-binding protein. J Biol Chem 2000; 275: 15498–503.

20. Guo S, Xu X, Ouyang Y, et al. Microarray expression profile analysis of circular RNAs in pancreatic cancer. Mol Med Rep 2018; 17: 7661–7671.

21. Li H, Hao X, Wang H, et al. Circular RNA Expression Profile of Pancreatic Ductal Adenocarcinoma Revealed by Microarray. Cell Physiol Biochem 2016; 40: 1334–1344.

22. Chen G, Shi Y, Zhang Y, et al. CircRNA_100782 regulates pancreatic carcinoma proliferation through the IL6-STAT3 pathway. Onco Targets Ther 2017; 10: 5783–5794.

23. Zhu P, Ge N, Liu D, et al. Preliminary investigation of the function of hsa_circ_0006215 in pancreatic cancer. Oncol Lett 2018; 16: 603–611.

24. Li Z, Yanfang W, Li J, et al. Tumor-released exosomal circular RNA PDE8A promotes invasive growth via the miR-338/MACC1/MET pathway in pancreatic cancer. Cancer Lett 2018; 432: 237–250.

25. Zhang J, Liu H, Hou L, et al. Circular RNA_LARP4 inhibits cell proliferation and invasion of gastric cancer by sponging miR-424-5p and regulating LATS1 expression. Mol Cancer 2017; 16: 151.

26. Yang C, Yuan W, Yang X, et al. Circular RNA circ-ITCH inhibits bladder cancer progression by sponging miR-17/miR-224 and regulating p21, PTEN expression. Mol Cancer 2018; 17: 19.

27. Liu L, Liu FB, Huang M, et al. Circular RNA ciRS-7 promotes the proliferation and metastasis of pancreatic cancer by regulating miR-7-mediated EGFR/STAT3 signaling pathway. Hepatobiliary Pancreat Dis Int 2019; pii: 30039–6.

28. An Y, Cai H, Zhang Y, et al. circZMYM2 Competed Endogenously with miR-335-5p to Regulate JMJD2C in Pancreatic Cancer. Cell Physiol Biochem 2018; 51: 2224–2236.

29. Cao Z, Liu C, Xu J, et al. Plasma microRNA panels to diagnose pancreatic cancer: Results from a multicenter study. Oncotarget 2016; 7:41575–41583.

30. Omura N, Mizuma M, MacGregor A, et al. Overexpression of ankyrin1 promotes pancreatic cancer cell growth. Oncotarget 2016; 7: 34977–87.

31. Okada Y, Eibl G, Duffy JP, et al. Glial cell-derived neurotrophic factor upregulates the expression and activation of matrix metalloproteinase-9 in human pancreatic cancer. Surgery 2003; 134: 293–9.

32. Mascarenhas JB, Young KP, Littlejohn EL, et al. PAX6 is expressed in pancreatic cancer and actively participates in cancer progression through activation of the MET tyrosine kinase receptor gene. J Biol Chem 2006; 284: 27524–32.

33. Yang Q, Du WW, Wu N, et al. A circular RNA promotes tumorigenesis by inducing c-myc nuclear translocation. Cell Death Differ 2017; 24: 1609–1620.

34. Yang F, Fang E, Mei H, et al. Cis-Acting circ-CTNNB1 Promotes β-Catenin Signaling and Cancer Progression via DDX3-Mediated Transactivation of YY1. Cancer Res 2019; 79: 557–571.

35. Heumann A, Kaya Ö, Burdelski C, et al. Up regulation and nuclear translocation of Y-box binding protein 1 (YB-1) is linked to poor prognosis in ERG-negative prostate cancer. Sci Rep 2017; 7: 2056.

36. Lim JP, Nair S, Shyamasundar S, et al. Silencing Y-box binding protein-1 inhibits triple-negative breast cancer cell invasiveness via regulation of MMP1 and beta-catenin expression. Cancer Lett 2019l; 452: 119–131.

37. Chua PJ, Lim JP, Guo TT, et al. Y-box binding protein-1 and STAT3 independently regulate ATP-binding cassette transporters in the chemoresistance of gastric cancer cells. Int J Oncol 2018; 53: 2579–2589.

38. Lu J, Li X, Wang F, et al. YB-1 expression promotes pancreatic cancer metastasis that is inhibited by microRNA-216a. Exp Cell Res 2017; 359: 319–326.

39. Gallardo M, Hornbaker MJ, Zhang X, et al. Aberrant hnRNP K expression: All roads lead to cancer. Cell Cycle 2016; 15: 1552–7.

40. Liu XH, Ma J, Feng JX, et al. Regulation and related mechanism of GSN mRNA level by hnRNPK in lung adenocarcinoma cells. Biol Chem 2019; 400: 951–963.

41. Gao R, Yu Y, Inoue A, et al. Heterogeneous nuclear ribonucleoprotein K (hnRNP-K) promotes tumor metastasis by induction of genes involved in extracellular matrix, cell movement, and angiogenesis. J Biol Chem 2013; 288: 15046–56.

42. Lu J, Gao FH. Role and molecular mechanism of heterogeneous nuclear ribonucleoprotein K in tumor development and progression. Biomed Rep 2016; 4: 657–663.

43. Hu P, Chen X, Sun J1, et al. siRNA-mediated knockdown against NUF2 suppresses pancreatic cancer proliferation in vitro and in vivo. Biosci Rep 2015; 35: pii: e00170.

