## Supplementary material for "CircFOXK2 Promotes Tumor Growth and Metastasis of Pancreatic Ductal Adenocarcinoma via Complexing with RNA Binding Proteins and Sponging MiR-942": suppl data

**Supplemental Information**

**Supplemental Methods**

**RNase R treatment**

5 μg of total RNA was digested with or without 15 U of RNase R (Epicentre) at 37 °C for 15 mins. RNA was then purified by TRIZOL Reagent.

**RNA stability assay**

PANC-1 cells were seeded to a 24-well plate. After 24 h, transcription was blocked by treating cells either 2 μg/mL actinomycin D (Sigma) or DMSO as a control for 0 h, 4 h, 12 h and 24 h. RNA was then purified by TRIZOL Reagent.

**Plasmid and oligonucleotide transfection**

circRNA overexpression plasmid was constructed by cloning the FOXK2 exon 2 and 3 into pcDNA3.1 (+) circRNA mini vector (Addgene). pmiR-Reporter plasmid with the inclusion of circRNA sequence in the 3’UTR for luciferase assay was constructed by cloning circRNA sequence into region directly downstream of the firefly luciferase gene in the pmiR-Reporter (Promega). Mutation of the miRNA binding site in the pmiR-Reporter plasmid was generated using KAPA HiFi DNA Polymerase (KapaBiosystem) and primers with mutation sites. Lentiviral vector for stable knockdown of circFOXK2 was generated by cloning the shRNA sequence targeting circFOXK2 into lentiviral vector with H1 promoter as described previously^16^. miRNAs and siRNAs were purchased from GenePharma. Plasmids, miRNAs and siRNAs transfection were performed by lipofectamine 3000 (Invitrogen), according to the manufacturer’s protocol.

**Lentiviral production and infection**

The VSV-G-pseudotyped lentivirus was produced by co-transfecting packaging vectors: pCMV-VSVG, pRSV-REV and pMDLg/pRRE with transfer vectors in HEK293T cells as described previously^17^. For lentiviral infection, 2,500 cells were seeded in a 24-well plate. After 24 h, cells were infected with lentivirus with polybrene for 72 h. Transfection medium was changed and cells were cultured with 800 μg/ml geneticin for 2 weeks. The efficiency of knockdown was confirmed by qRT-PCR.

**Quantitative reverse transcription PCR (qRT-PCR)**

The cytoplasmic and nuclear fractions were extracted using NE-PER Nuclear and Cytoplasmic Extraction Reagents (Thermo Fisher Scientific). RNA from whole-cell lysate or cell fractions were isolated by TRIZOL Reagent. Formalin-fixed, paraffin-embedded (FFPE) sample RNA was isolated by miRNeasy FFPE Kit (Qiagen) according to manufacturer’s protocol. Measurement of gene expression level was performed by qRT-PCR. Reverse transcription of total RNA (except miRNA) was performed by High-Capacity cDNA Reverse Transcription Kit (Applied Biosystems). Reverse transcription of miRNA was performed by Mir-X™ miRNA First-Strand Synthesis Kit (Takara). qRT-PCR was performed by ABI 7900HT Real-Time PCR system using SYBR Green PCR Master Mix (Applied Biosystems). The primers used in this study were listed in supplemental table 1.

**Cell viability assay**

3-(4,5-dimethylthiazol-2-yl)-2,5-diphenyltetrazolium bromide (MTT) cell viability assay was performed by seeding 2500 cells in a 96-well plate. After 24, 72 and 120 h, the medium was removed and incubated with 0.65 mg/mL MTT in DMEM. After 2 h, the mixture was removed, and the insoluble formazan was solubilized by adding 120 μL DMSO. After 5 mins, absorbance at 570 nm was measured using microplate spectrophotometer (Biorad).

**Anchorage-dependent colony formation assay**

Anchorage-dependent colony formation assay was performed by seeding 1000 cells in a 6-well plate. After growing the cells for 2 weeks, the cells were stained by 0.5% crystal violet. After staining for 5 mins, the cells were washed with PBS to reduce background. Photos were taken and number of colonies was counted.

**Anchorage-independent colony formation assay**

A base agar was prepared by mixing 1.2% agar solution with 2X culture medium and was allowed to solidify in a 6-well plate. The 5000 cells in 2X culture medium was mixed with 0.7% agar solution and was added on the top of solidified base agar. When the top layer of agar with cells was solidified. 1 mL of culture medium was added to the top agar layer. After growing cells for 3 weeks, the cells were stained with 0.05% crystal violet. After staining for 1 h, the cells were washed with PBS to reduce background. Photos were taken and number of colonies was counted.

**Cell migration assay**

Wound healing cell migration assay was performed by seeding 500000 cell/ mL in a 3-well silicone insert with a defined cell-free gap in a 24-well plate. On the next day, in which cells reached 90% confluency, the insert and medium were removed and culture medium without serum was added. Wound width was measured at 0, 24 and 48 h.

**Cell invasion assay**

Upper chamber of the trans-well insert with pore size of was coat with Matrigel (Corning) and was placed in a 37°C incubator overnight. Then the unsolidified Matrigel was removed and was seeded with 25000 cells in culture medium without serum. Then the insert with seeded cells was placed in a plate well with complete culture medium. After 24 to 72 h, the trans-well was fixed with 3.7 % formaldehyde for 5 mins, permeabilized by methanol for 15 mins, and was then stained by 0.5 % crystal violet for 20 mins. The cells in the upper chamber (non-invasive cells) were removed and the invaded cells were counted.

**Cell cycle analysis**

After knockdown of circRNAs for 72 h, cells were trypsinized and washed twice with 2 % FBS in PBS. Cells were then fixed with 70 % ethanol at 4 °C for 3 h. After ethanol fixation, cells were washed twice with 2 % FBS in PBS, followed by treatment with 50 μg/mL RNase A (Thermo Fisher Scientific) and 10 μg/mL propidium iodide at 37 °C for 15 mins. Cell cycle was analyzed by BD LSR Fortessa Cell Analyzer (BD Biosciences, USA). The cell cycle phase distribution and proportion of apoptotic cells were analyzed using BD FACSDiva software.

**Apoptosis assay**

Cell apoptosis assay after knockdown of circRNAs was performed by Annexin V-Cy5 Apoptosis Detection Kit (BioVision). Cells in 24-well plate were washed twice with ice-cold PBS. Then cells were stained by DAPI and 5 μL Annexin V in 500 μL 1X Annexin V Binding Buffer. After incubation at room temperature for 5 mins in dark, apoptotic cells were analyzed under fluorescent microscope.

**Immunoblotting**

Whole cell extract was prepared by lysing cells in NP-40 lysis buffer with proteinase inhibitors (Roche) and phosphatase inhibitor (Pierce). Protein concentration was determined by BCA assay (Pierce). Proteins were resolved by SDS-PAGE at different percentages, transferred to PVDF membrane and immunoblotted overnight at 4 °C with antibodies against PAX6 (rabbit; Cell Signaling #60433; 1:1000); ANK1 (rabbit; abcam ab58698; 1:1000); GDNF (rabbit; abcam ab18956; 1:1000); Caspase 3 (rabbit; Cell Signaling #9665 ;1:1000); PARP (rabbit; Cell Signaling #9542; 1:1000); Bcl-2 (rabbit; millipore 04-436; 1:1000); and GAPDH (rabbit; Cell signaling 5174; 1:1000). Then, the blots were washed three times with TBST, followed by incubation with 1:2000 secondary anti-rabbit antibody at room temperature for 1 h. Chemiluminescent signals were developed using Clarity™ Western ECL Substrate (Bio-Rad). Images of immunoblot were taken with ChemiDoc Imaging System (Bio-Rad).

**Immunohistochemical Staining**

Immunohistochemical staining was performed using human FFPE PDAC tumor samples. The sectioned tissues were deparaffinized and rehydrated by xylene and a series of graded ethanol. Antigen retrieval was performed using PT module (Thermofisher). Then, IHC was performed using Histostain-Plus IHC Kit, HRP, broad spectrum (Life Technologies, Carlsbad, CA). The sections were probed with appropriate antibody. Sections were counter-stained with hematoxylin, were dehydrated with a series of graded ethanol and xylene, and were mounted. A scoring system, based on the percentage of positive cells and staining intensity under the microscope with 100X magnification, was used to quantify the staining. 4 categories (0, 1, 2, and 3) were demoted as 0%, 1-10%, 10-50%, and >50%.

**Supplemental Figure**

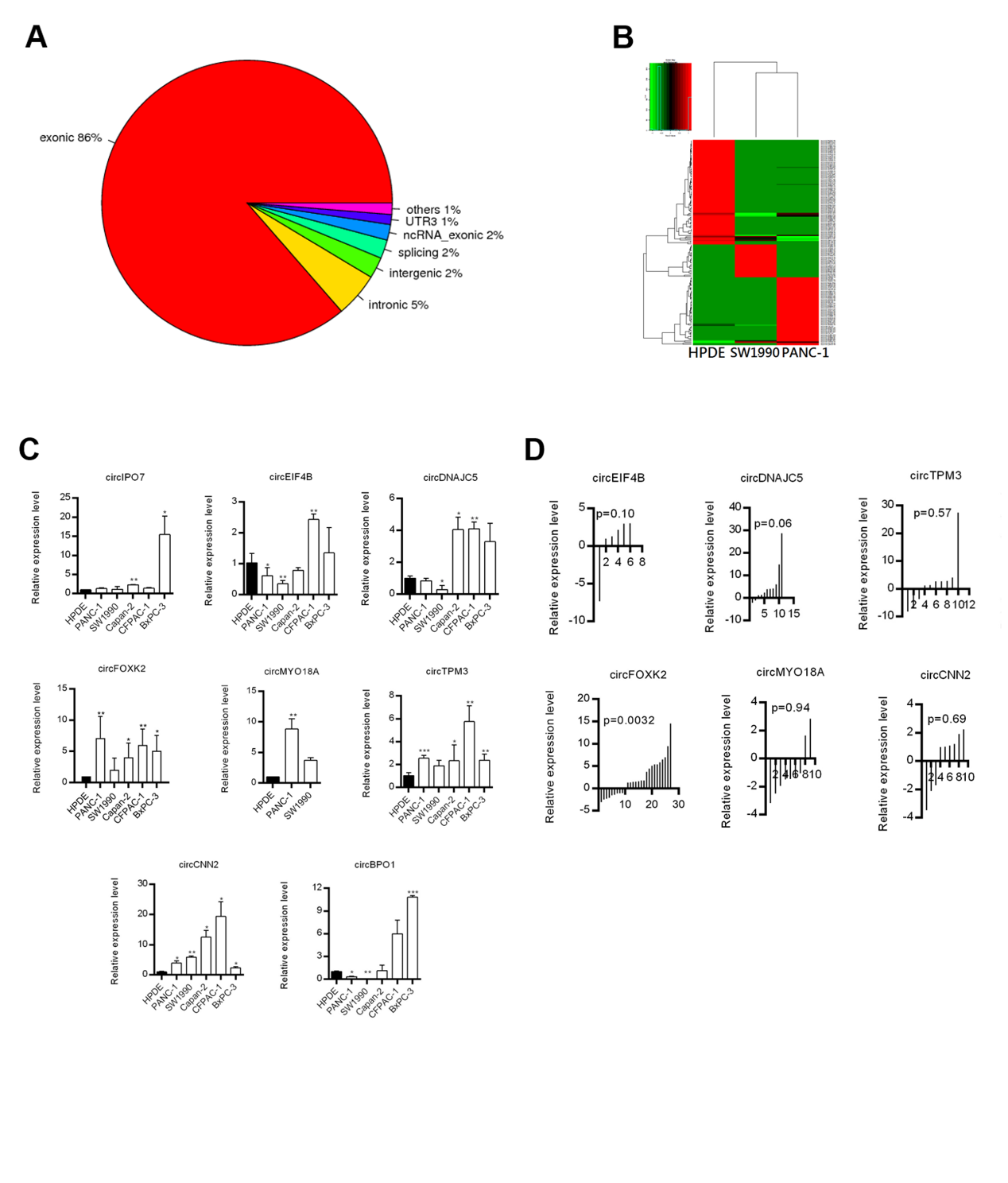

**Supplemental figure 1.** Identification of differentially expressed circRNAs in PDAC.

A, Distribution of circRNAs in the gene locations. The genomic location of each circRNA identified by circRNA sequencing was analyzed.

B, Clustered heatmap for circRNAs from normal HPDE and PDAC cells.

C, circRNA profiling in PDAC cell lines. The expression level was compared to normal HPDE cells.

D, circRNA profiling in PDAC tumors. The expression level of circRNAs in tumor sample was compared to respective adjacent non-tumor tissue.

Data are from at least three independent experiments Mean ± SD (*p<0.05; **p<0.01; ***p<0.001)

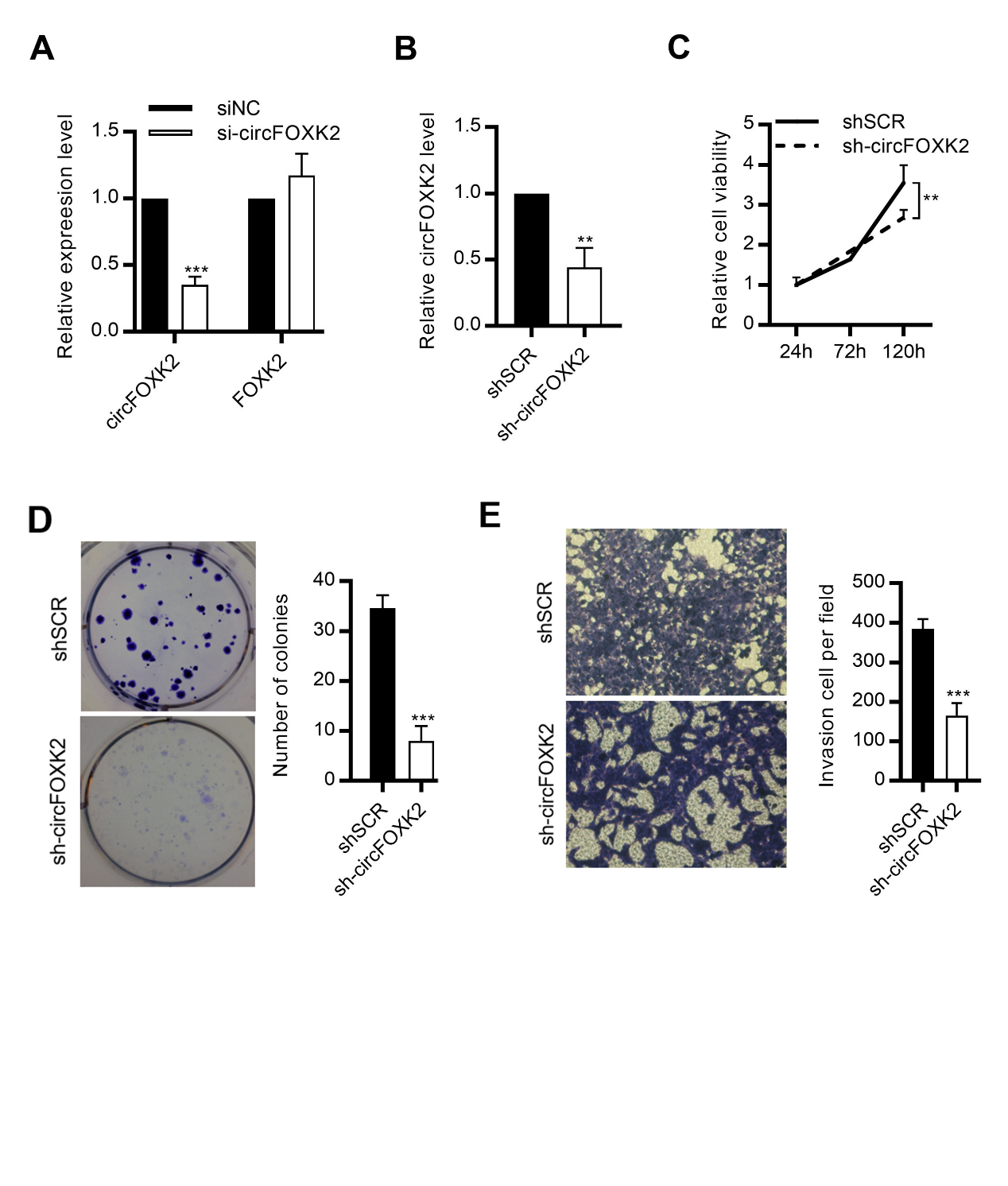

**Supplemental figure 2.** circFOXK2 promotes PDAC cell growth and invasion.

A, Knockdown efficiency of circFOXK2 by siRNAs. The expression levels of circFOXK2 and FOXK2 were analyzed by qRT-PCR.

B, Knockdown efficiency of circFOXK2 by shRNAs.

C, Cell growth analyzed by MTT assay after stable knockdown of circFOXK2 by shRNA for 24, 72 and 120 h in CFPAC-1 cells.

D, Anchorage-dependent colony formation after stable knockdown of circFOXK2 in CFPAC-1 cells. Cells were stained with crystal violet.

E, Trans-well cell invasion assay after stable knockdown of circFOXK2 in CFAPC-1 cells. Cells were stained by crystal violet.

Data are from at least three independent experiments Mean ± SD (**p<0.01; ***p<0.001)

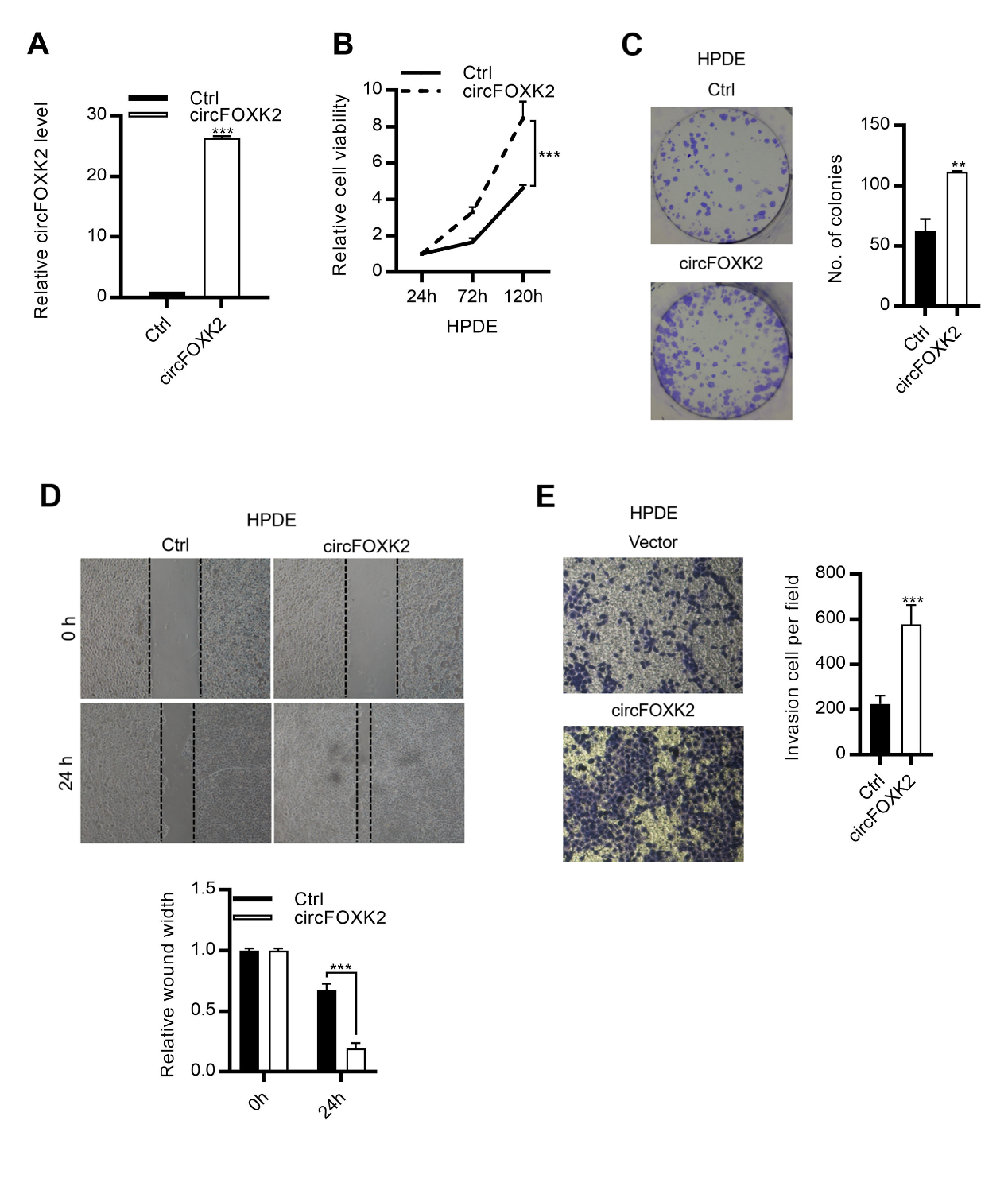

**Supplemental figure 3.** circFOXK2 promotes cell growth and invasion.

A, Overexpression efficiency of circFOXK2 in HPDE cells.

B, Cell growth analyzed by MTT assay after overexpression of circFOXK2 for 24, 72 and 120 h.

C, Anchorage-dependent colony formation after overexpression of circFOXK2. Cells were stained with crystal violet.

D-E (D) Wound healing cell migration assay and (E) trans-well cell invasion assay after overexpression of circFOXK2. Cells in invasion assay were stained by crystal violet.

Data are from at least three independent experiments Mean ± SD (**p<0.01; ***p<0.001)

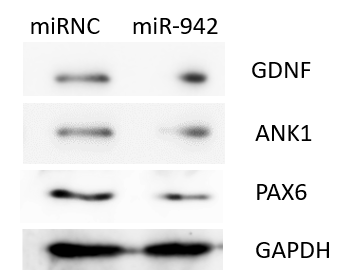

**Supplemental figure 4.** Analysis of ANK1, GDNF and PAX6 levels after transfecting miR-942 mimics in PANC-1 cells.

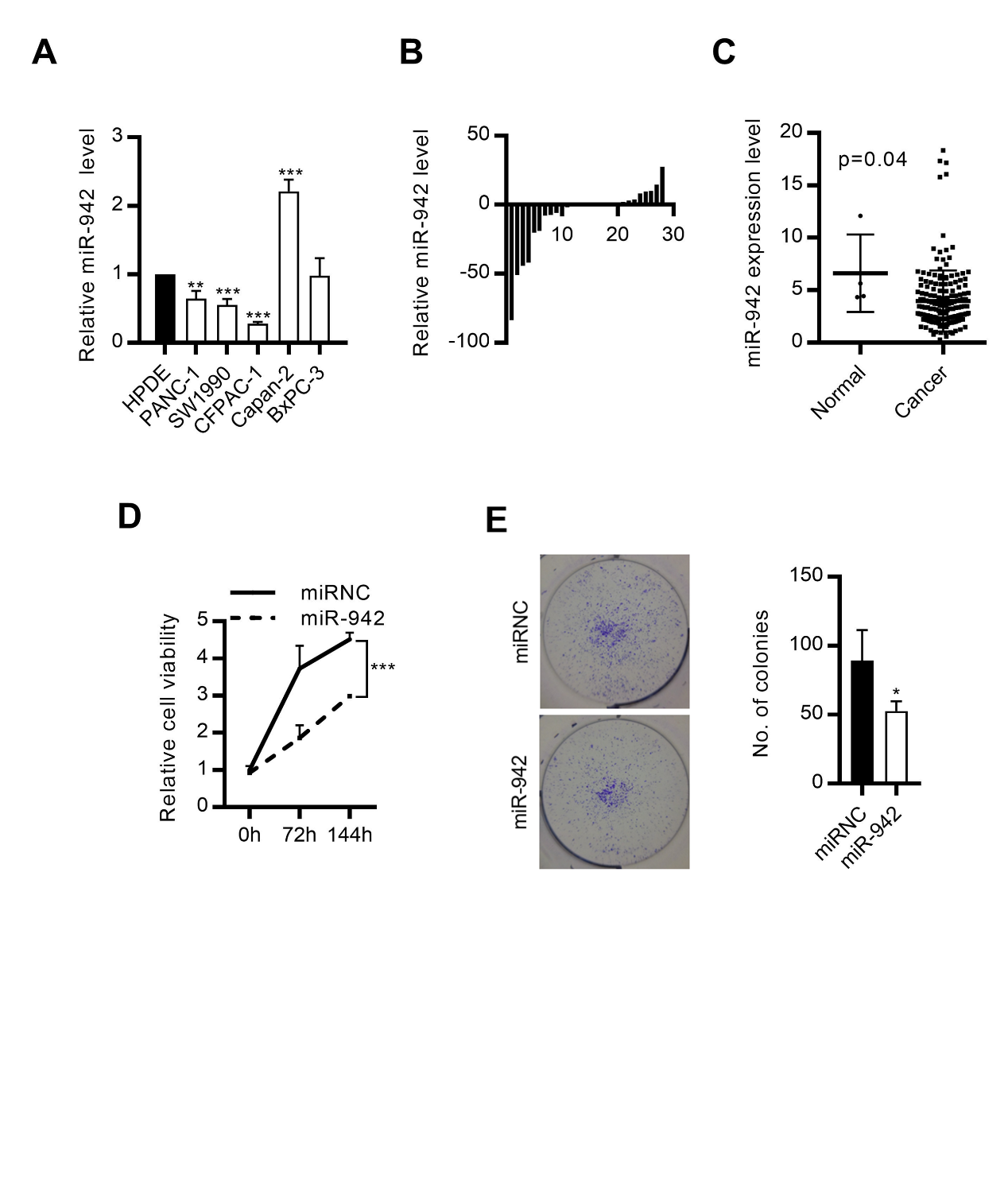

**Supplemental figure 5.** miR-942 inhibits PDAC development.

A-C Expression level of miR-942 in (A) PDAC cell lines, (B) tumors and (C) TCGA dataset.

D. Cell growth analyzed by MTT assay after transfecting miR-942 mimics for 24, 72 and 144 h in CFPAC-1 cells.

E. Anchorage-dependent colony formation after transfecting miR-942 mimics in CFPAC-1 cells. Cells were stained with crystal violet.

Data are from at least three independent experiments Mean ± SD (*p<0.05; **p<0.01; ***p<0.001)

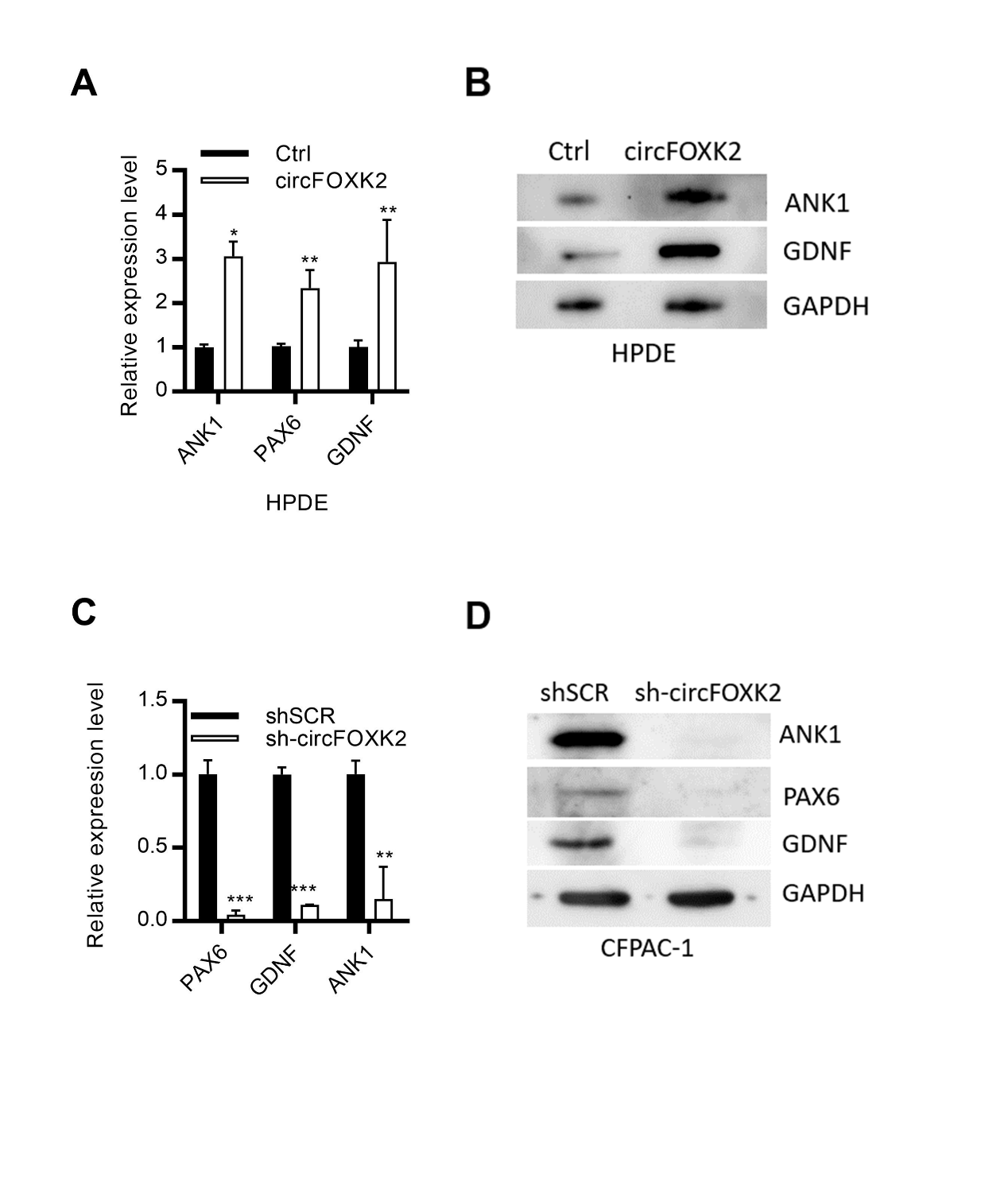

**Supplemental figure 6.** circFOXK2 promotes the expression of miR-942 targets in PDAC cells.

A-B. Analysis of ANK1, GDNF and PAX6 (A) mRNA and (B) protein levels after overexpressing circFOXK2 in HPDE cells.

C-D. Analysis of ANK1, GDNF and PAX6 (C) mRNA and (D) protein levels after stable knockdown of circFOXK2 in CFPAC-1 cells.

Data are from at least three independent experiments Mean ± SD (*p<0.05; **p<0.01; ***p<0.001)

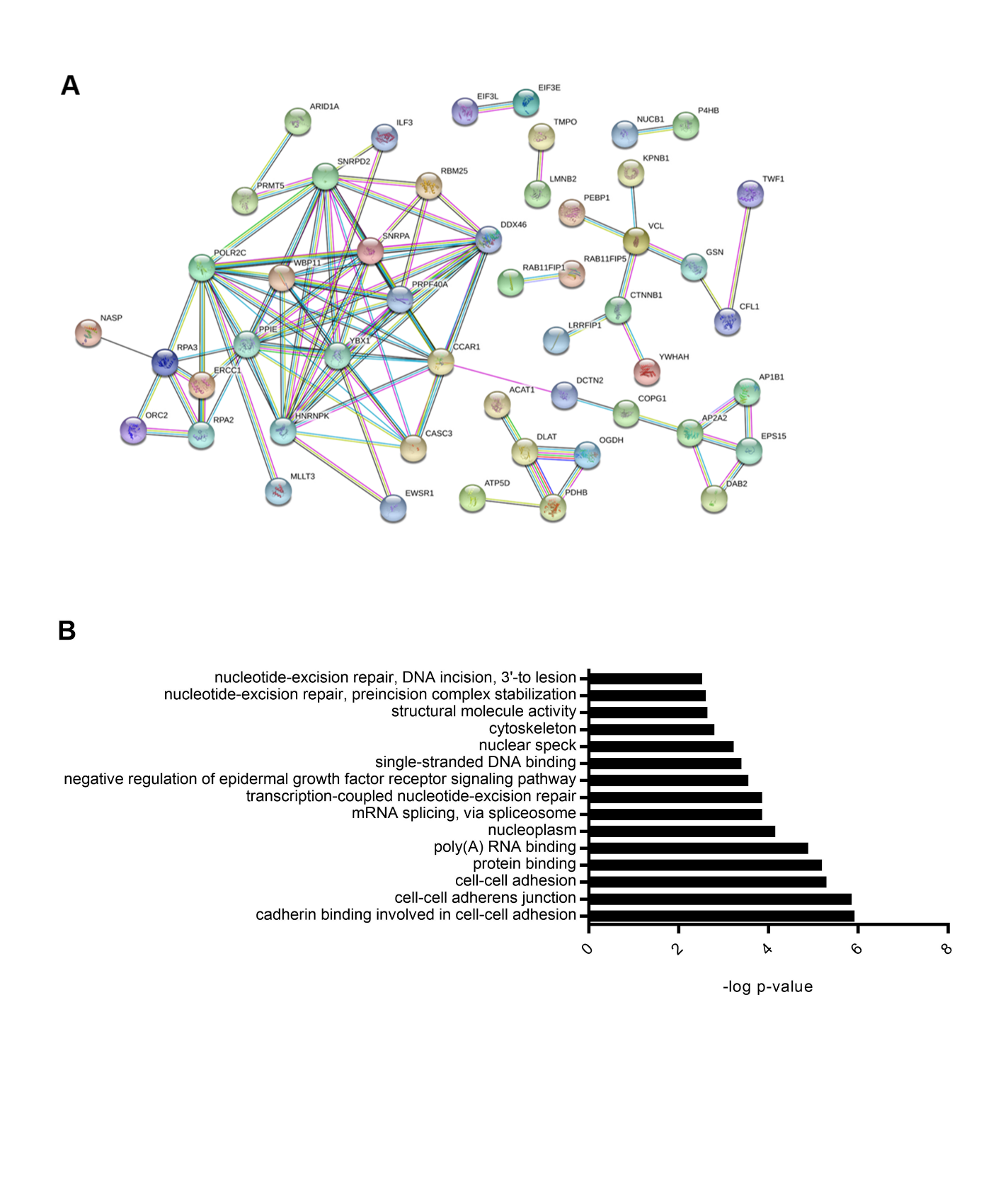
 **Supplemental figure 7.** Identification of circFOXK2-interacting proteins in PDAC.

A, Analysis of protein-protein interactions of circFOXK2-interacting proteins by STRING 11. Each node represents a protein and each edge represents a protein-protein interaction. Color of each edge denotes the type of evidence of the interaction. (Confidence level was set at high confidence of 0.7).

B, Gene Ontology analysis of the circFOXK2-interacting proteins in PDAC. The top 20 Gene Ontology clusters were listed.

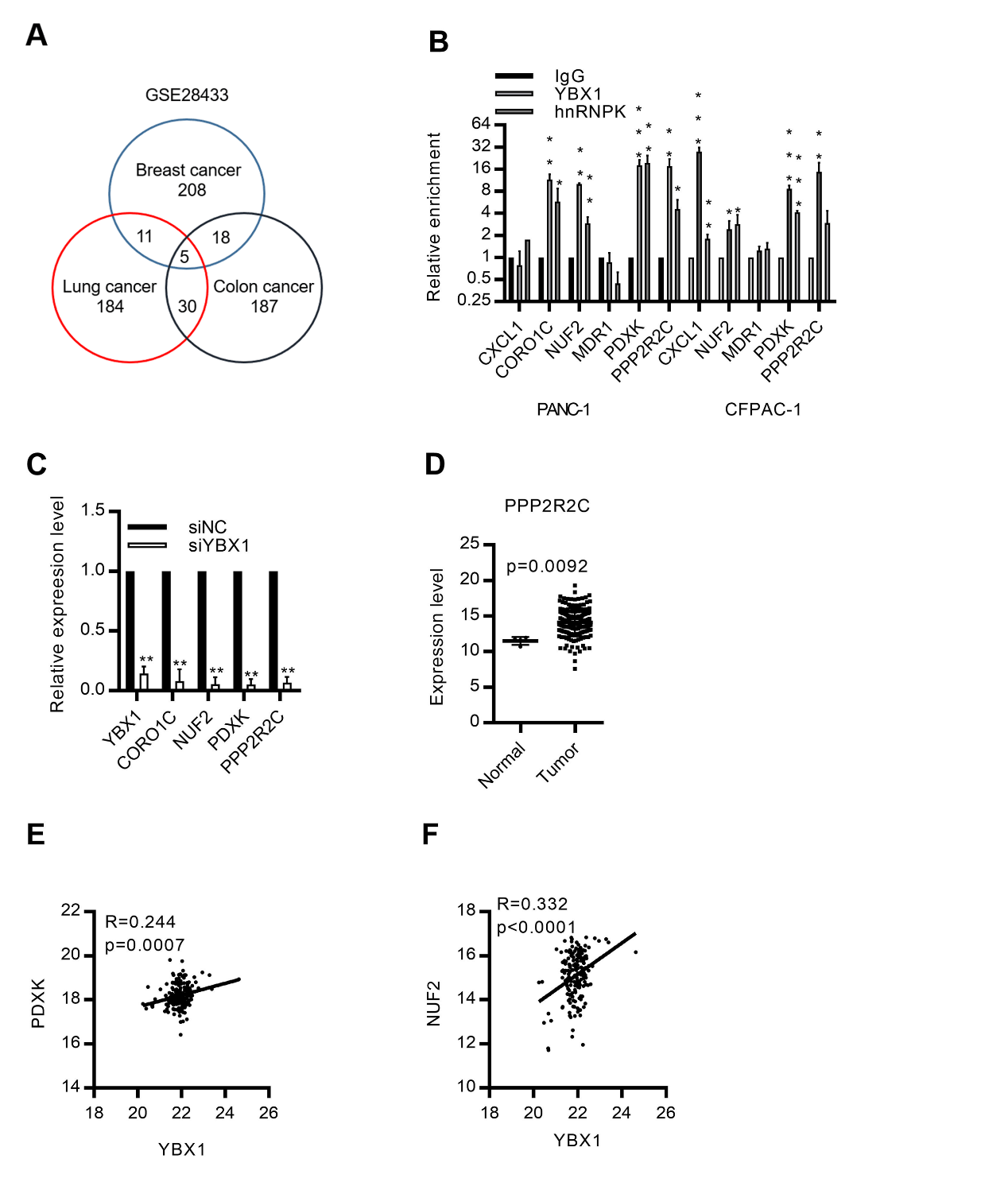
 **Supplemental figure 8.** Identification of YBX1/hnRNPK targets in PDAC.

A. Identification of YBX1 targets in multiple cancers.

B. RNA immunoprecipitation for YBX1 and hnRNPK in PDAC cells. The enrichment levels of YBX1 targets CXCL1, CORO1C, NUF2, MDR1, PDXK and PPP2R2C were analyzed by qRT-PCR.

C. Expression levels of YBX1 targets after knockdown of YBX1 in CFPAC-1 cells.

D-F. Analysis of (D) PPP2R2C, (E) PDXK and (F) NUF2 expression level and the correlation to YBX1 level in PDAC TCGA dataset.

Data are from at least three independent experiments Mean ± SD (*p<0.05; **p<0.01; ***p<0.001)

**Supplemental table 1. Primers used in this study**

| circFOXK2-F | CGGGGTCTTCAGGGTACAAGG |
| --- | --- |
| circFOXK2-R | GATGTTTGTGCTCGGGAACCT |
| FOXK2-F | GTGCACATTCAGGTTCCCGAG |
| FOXK2-R | CTTCGGGCTGTCTCCACCTGA |
| PAX6-F | ACATTTCTGCAGGGGAGTGA |
| PAX6-R | TTACAGCCAGCGAGAAGGAA |
| GDNF-F | AGCTGAGACAACGTACGACA |
| GDNF-R | GCCGGAGTCAGATACATCCA |
| ANK1-F | GGCCCGCAACGACGACACG |
| ANK1-R | AATGTTGCAGGGGCGTGAATC |
| YBX1-F | GGACAAGAAGGTCATCGCAAC |
| YBX1-R | TCTCCATCTCCTACACTGCGA |
| hnRNPK-F | CTGCTTCAGAGCAAGAATGCT |
| hnRNPK-R | AACTGCAGGCCCTCTTCCA |
| CABLES1-F | AGGGCCATTTCTTCTCAGCT |
| CABLES1-R | GCTACACACCCAAGACTCCT |
| NUF2-F | GCCGGGTGAATGACTTTGAG |
| NUF2-R | ACTGTTGCATTTTGTCCGCA |
| PDXK-F | GTGTGGCTGGACTGTACTCT |
| PDXK-R | GCACATAACCTGCTCTGCTC |
| PPP2R2C-F | TTTGAATGTGCCTGGAACGG |
| PPP2R2C-R | TCCAAGCTGTCCACACTGAT |
| CXCL1-F | AACCGAAGTCATAGCCACAC |
| CXCL1-R | GTTGGATTTGTCACTGTTCAGC |
| CORO1C-F | TCCTCCCTCTGCACAAGACT |
| CORO1C-R | GGATCTGCCATACCATGACC |
| U6-F | CGGCAGCACATATAC |
| U6-R | TTCACGAATTTGCGTGTCAT |
| GAPDH-F | TGCCTCCTGCACCACCAACT |
| GAPDH-R | CCCGTTCAGCTCAGGGATGA |
